# Microstructured hydrogels to guide self-assembly and function of lung alveolospheres

**DOI:** 10.1101/2021.08.30.457534

**Authors:** Claudia Loebel, Aaron I. Weiner, Jeremy B. Katzen, Michael P. Morley, Vikram Bala, Fabian L. Cardenas-Diaz, Matthew D. Davidson, Kazushige Shiraishi, Maria C. Basil, Matthias Ochs, Michael F. Beers, Edward E. Morrisey, Andrew E. Vaughan, Jason A. Burdick

**Affiliations:** Department of Bioengineering, University of Pennsylvania, Philadelphia, PA, USA; School of Veterinary Medicine, University of Pennsylvania, Philadelphia, PA, USA; Department of Medicine, Penn Center for Pulmonary Biology, University of Pennsylvania, Philadelphia, PA, USA; Department of Biomedical Engineering, University of Michigan, Ann Arbor, MI, USA; Institute of Functional Anatomy, Charité - Universitätsmedizin Berlin, Berlin, Germany

**Author notes:** Department of Materials Science & Engineering, University of Michigan, Ann Arbor, MI, USA.

## Abstract

Epithelial cell organoids have increased opportunities to probe questions on tissue development and disease *in vitro* and for therapeutic cell transplantation. Despite their potential, current protocols to grow these organoids almost exclusively depend on culture within three-dimensional (3D) Matrigel, which limits defined culture conditions, introduces animal components, and results in heterogenous organoids (i.e., shape, size, composition). Here, we describe a method that relies on polymeric hydrogel substrates for the generation and expansion of lung alveolar organoids (alveolospheres). Using synthetic hydrogels with defined chemical and physical properties, human induced pluripotent stem cell (iPSC)-derived alveolar type 2 cells (iAT2s) self-assemble into alveolospheres and propagate in Matrigel-free conditions. By engineering pre-defined microcavities within these hydrogels, the heterogeneity of alveolosphere size and structure was reduced when compared to 3D culture while maintaining alveolar type 2 cell fate of human iAT2 and primary mouse tissue-derived progenitor cells. This hydrogel system is a facile and accessible culture system for the culture of primary and iPSC-derived lung progenitors and the method could be expanded to the culture of other epithelial progenitor and stem cell aggregates.

## Main

Organoids have received considerable attention for modeling organogenesis and disease, facilitating large screening of therapeutic molecules (e.g., proteins, drugs), and for the sourcing of cells for therapeutic transplantation^1,2^. With regards to the lung, organoids have primarily been applied to address questions in pulmonary biology, such as reparative mechanisms of lung progenitors and their responsiveness to regenerative and therapeutic molecules^3^. Given the importance of gas-exchange as the most fundamental function of the lungs, alveolar organoids (i.e., alveolospheres) are becoming an indispensable tool for *in vitro* studies, including for the modeling of distal lung injuries such as SARS-CoV-2 infections^4–6^. The alveolar epithelium comprises two distinct epithelial cell types: Type 2 cells (AT2), which are surfactant -producing alveolar progenitor cells that can self-renew and differentiate into type 1 cells (AT1) that cover the majority of the surface of the lung alveoli^7,8^. When cultured in Matrigel and with mesenchymal feeder cells, primary mouse-derived AT2 cells form alveolospheres that comprise both AT1 and AT2 cells but often do not reproduce the structure of alveoli in adult lungs^3^. Importantly, recent advances in the differentiation of induced pluripotent stem cells (iPSCs) into AT2 cells have also enabled the culture of human alveolospheres, overcoming some of the challenges in the isolation and culture of primary human AT2 cells^9^.

Although the spontaneous assembly and propagation of organoids within Matrigel have resulted in significant advances, artificial niches are increasingly being developed to provide distinct chemical and physical signals to organoids during culture^10,11^. Specifically, synthetic niches can guide cellular self-assembly while overcoming challenges in Matrigel organoid cultures, such as the inherent variability in organoid formation efficiencies and morphology, which poses issues for standardizing organoid models across laboratories with quantitative readouts^1,2,12^. Several engineering modalities such as customized synthetic hydrogels^13–16^, bioprinting^17,18^ and microfluidics^19,20^ have been implemented to increase the number of controllable parameters during organoid cultures, such as cell-matrix interactions, the organization of multiple cell types, and fluid flow. Recent efforts have also focused on matrix-free approaches, including the design of polymeric substrates with cavities to provide geometrical constraints for the generation of intestinal and pancreatic organoids^21,22^ ^23^. However, little is known of how geometrical cues of matrix-free culture conditions might guide the generation and function of lung alveolospheres.

Here, we propose hydrogel culture systems as methods to form and expand lung alveolospheres. First, encapsulation within 3D hydrogels was pursued with synthetic hyaluronic acid (HA) hydrogels, which allowed for alveolosphere formation and growth when modified with laminin/entactin, but with similar heterogeneity to Matrigel controls. As an alternative, a microstructured hydrogel was developed to guide lung alveolosphere formation within microwells under Matrigel-free culture conditions. Building upon recent advances in the design of defined matrices for organoids, our approach uses synthetic HA hydrogels that are engineered to contain pre-defined microcavities to generate uniform lung alveolospheres within individual microwells. Control of the initial aggregate size through cell seeding densities and microwell size enabled us to explore the role of culture constraints on alveolosphere growth and maturation. Using this method, we generated human iPSC-derived alveolospheres that display AT2 functional capacities, such as expression and processing of surfactant proteins. Alveolospheres were also generated from primary mouse alveolar progenitors, providing a minimally engineered and Matrigel-free approach for guided lung organoid culture.

## Results

### Defined 3D hydrogels enable SFTPC^GFP+^ alveolosphere formation

To illustrate the potential of hydrogels to enable assembly of alveolosphere organoids, we first employed human iPSC-derived alveolar progenitor cells (iAT2s) with a GFP knock-in cassette in one allele of the SFTPC gene, a specific marker for AT2 cells (Fig. 1A)^24^. Using a recently developed protocol^9,25^, we differentiated iPSCs (RUES2 line) into foregut endoderm followed by sorting for NKX2.1+ putative lung primordial lung progenitors and consecutive enrichment for SFTPC^GFP+^ cells (Fig. 1A). The differentiation protocol resulted in a total yield of ∼55% SFTPC^GFP+^ cells on day 66 in Matrigel (Fig. 1B). The resulting SFTPC^GFP+^ cells were further purified and passaged as alveolospheres in the presence of lung maturation additives and selective Rho-associated kinase (ROCK) inhibitor, Y-27632 from day 0-2. CHIR was added back following 7 days of withdrawal (days 2-9) to increase efficiency of iAT2 maturation^9^ (Fig. 1C). Thus, this protocol allows for iAT2 serial passaging upon dissociation into single cells and formation of alveolospheres that retain SFTPC^GFP+^ expression in Matrigel.

**Figure 1.**
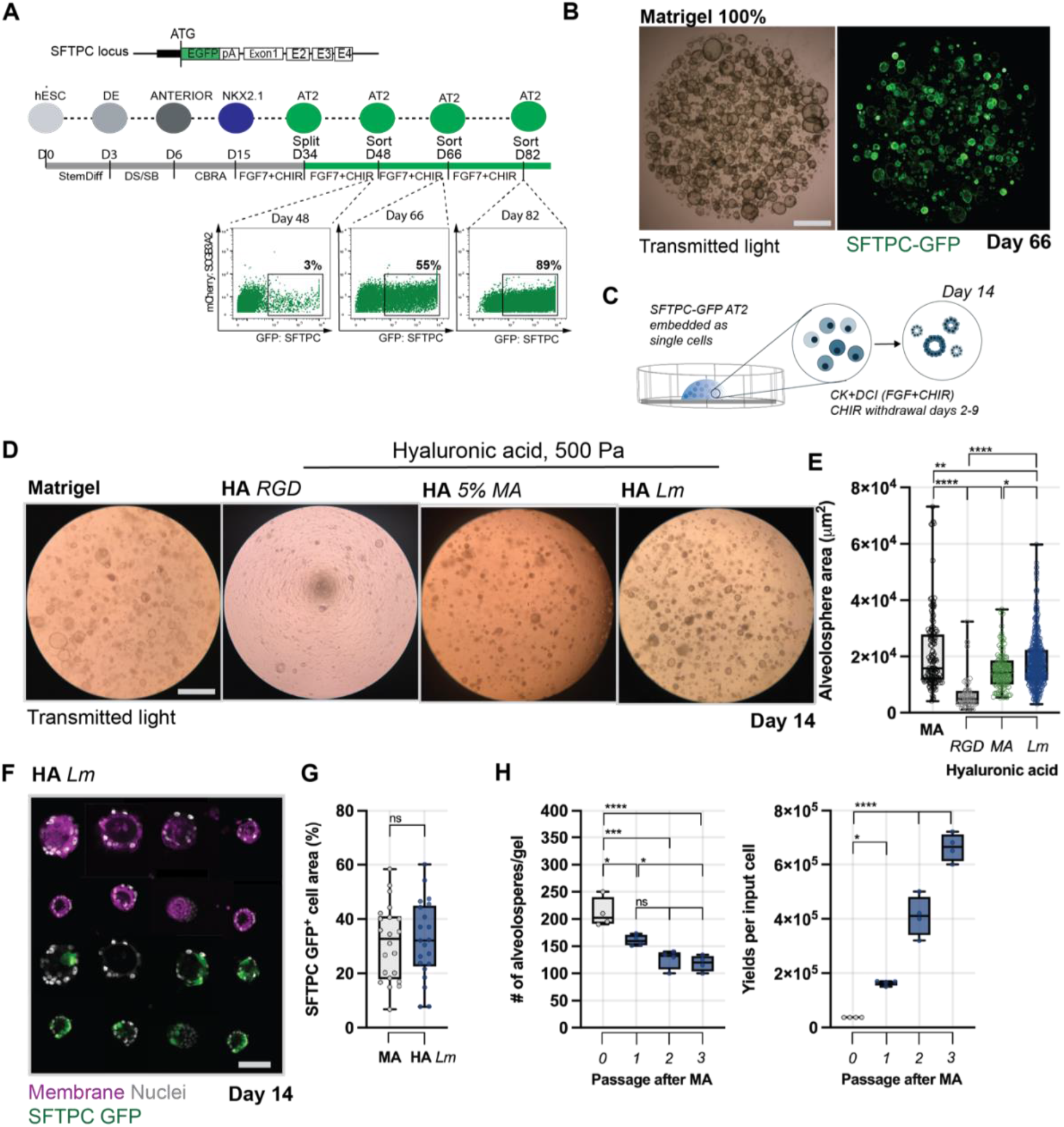
Tailored 3D hydrogel matrices for alveolosphere assembly and growth. **A** Edited SFTPC^GFP+^ loci post Cre-mediated antibiotic cassette excision and differentiation protocol from human ESCs (RUES2) to putative iAT2s with representative flow cytometry and enrichment of GFP^+^ cells used for subsequent studies. **B** Representative images of alveolospheres showing expression of SFTPC^GFP+^ at day 66 upon GFP^+^ enrichment and culture in Matrigel (scale bar 100 µm). **C** Schematic showing SFTPC^GFP+^ cells embedded into 3D hydrogels as a single cell suspension with 500 cells per µL and cultured with CHIR withdrawal between day 3-8 and CHIR addback (days 9-14). **D** Representative images of alveolospheres formed in Matrigel and hyaluronic acid hydrogels at 14 days. Hyaluronic acid hydrogels were crosslinked with a protease sensitive crosslinker via visible light-initiated thiol-ene reaction either modified with a cell-adhesive peptide (1 mM RGD), supplemented with 5% (wt/vol) Matrigel or 2 mg/mL laminin/entactin and with crosslinker amount adjusted to achieve an initial storage (elastic) modulus of 500 Pa (scale bar 100 µm, see supplementary figure 1 for representative images and quantification of viability and projected alveolosphere area in hyaluronic acid hydrogels supplemented with various concentrations of Matrigel, laminin/entactin and the influence of elastic modulus and cell seeding density). **E** Quantification of projected alveolosphere area at 14 days measured from brightfield images (*p<0.05, ****p<0.0001 by ANOVA and Bonferroni’s multiple comparisons test, n = 155 (Matrigel), n = 55 (HA RGD), n = 106 (HA Ma), and n = 154 (HA Lm)). **F** Representative images of SFTPC^GFP+^ alveolospheres in HA LM hydrogels at 14 days (cell mask membrane stain (magenta), nuclei (grey), and SFTPC^GFP+^ expression (green), scale bar 100 µm). **G** Quantification of SFTPC^GFP+^ per alveolosphere area (cell membrane stain) at 14 days (ns = not significantly different by unpaired two-tailed *t*-test). **H** Quantification of the number of alveolospheres and proliferation kinetics of cell yield per re-embedded SFTPC^GFP+^ cells in HA Lm hydrogels over three passages.

To demonstrate alveolosphere formation in synthetic hydrogels, iAT2s were embedded within hyaluronic acid (HA) hydrogels (norbornene-modified HA, modified with cell-adhesive peptide (HA *RGD*), 5% Matrigel (HA 5% MA) or 2 mg/mL Laminin/Entactin (HA *Lm*), all with a storage modulus of ∼500 Pa (Fig. 1D). At 14 days, small alveolospheres were observed in the HA *RGD* hydrogel, whereas culture in HA *MA* and HA *Lm* hydrogels resulted in increased formation of alveolospheres; however, the area was lower than MA controls (Fig. 1E). Alveolosphere formation further depended on the concentration of Matrigel and Laminin/Entactin, initial storage modulus and iAT2 seeding density (Supplementary Fig. 1), indicating that the formation of alveolospheres is influenced by both chemical and mechanical signals.

Alveolospheres formed within HA *Lm* hydrogels and expressed SFTPC^GFP+^ (Fig. 1F), with a colony forming efficiency that was similar to Matrigel (Supplementary Fig. 2). Furthermore, quantification of SFTPC^GFP+^ expression confirmed that alveolospheres maintain their AT2 progeny with an average of ∼31% SFTPC^GFP+^ cells (Fig. 1G). Given that HA *Lm* hydrogels support alveolosphere growth, we next assessed whether iAT2s retain proliferative potential during serial passaging, consistent with previous protocols in Matrigel^25^, but using a modified enzymatic digestion (1 mg/mL hyaluronidase, no dispase). Using hyaluronidase digestion and subsequent trypsin digestion into single cell suspensions, over a period of 3 passages, iAT2s reformed spheres (Supplementary Fig. 3) with stable efficiency and proliferation kinetics (Fig. 1H). These findings indicate that iAT2-derived alveolospheres can be formed and maintained in laminin/entactin-enriched synthetic hydrogels, enabling culture in a well-defined Matrigel-free 3D environment. However, this strategy presents relatively high heterogeneity, because alveolospheres form randomly, similar to traditional Matrigel culture. Thus, although well-defined HA hydrogels improve culture conditions, limited control over size and shape and difficulties of standardizing towards downstream analysis remain to be addressed.

### Design of microwell hydrogels for generation of alveolospheres

Given that iAT2s were able to form alveolospheres within a Matrigel-free hydrogel, we next sought to further control the growth and homogeneity of formed alveolospheres, a critical requirement for downstream read-outs. Thus, we engineered hydrogel substrates that comprise microwell-shaped cavities for aggregation and individual alveolosphere formation through geometrical constraints (Fig. 2). Accessibility and customization of such engineered systems is often the bottleneck for broader applications within the regenerative biology community. Here, we used commercially available cell culture surfaces (EZSPHERE^TM^) with evenly spaced microwells to make silicone replica molds of a desired size and shape (Supplementary Fig. 4). HA hydrogels were then fabricated with the silicone molds upon ultraviolet light mediated photo-curing onto a glass coverslip (Fig. 2A, i) and subsequent transfer of the fabricated microwell hydrogels directly onto conventional tissue culture dishes (Fig. 2A, ii). Using this fabrication technique, hydrogels can be readily fabricated and are compatible with a range of different sizes as shown by fluorescent images of individual microwells (Fig. 2B). As the microwell size can be accurately modulated, we fabricated hydrogels (∼20 kPa) of small, medium and large microwell sizes, ranging from an average of 80-300 µm in depth (amplitude) and 250-810 µm in width (Fig. 2C). Hydrogel structures were stable during 14 days in culture in alveolosphere media with minimal changes in amplitude and width of individual microwells (Supplementary Fig. 5).

**Figure 2.**
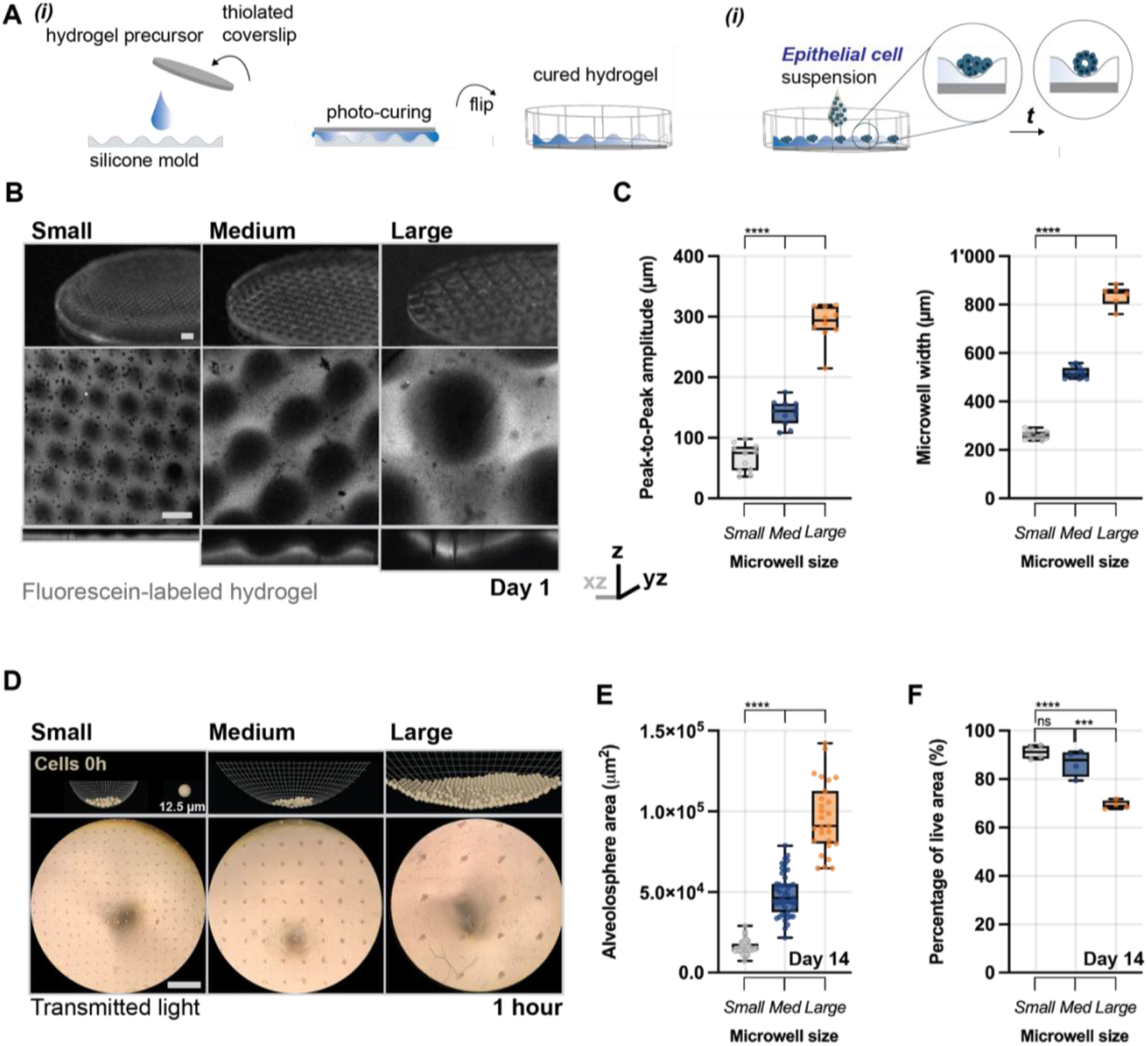
Design and characterization of microwell hydrogels for alveolosphere culture. **A** Schematic illustrating the fabrication of microwell hydrogels: (i) Silicone molds were generated by moulding EZSPHERE^TM^ microwell-shaped culture dishes. The microwells were then imprinted onto hydrogel surfaces during ultra-violet light-initiated crosslinking of a norbornene-modified hyaluronic acid hydrogel precursor solution. (ii) Single cells were seeded on top of the microwells in CD+CKI media and left to settle and form alveolospheres within individual microwells. **B** Representative images of microwell hyaluronic acid hydrogels modified with fluoresceine and fabricated from silicone molds with different widths and depths upon 1 day of swelling in saline (small: 200 µm/100 µm, medium (Med): 500 µm/200 µm, large: 800 µm/300 µm (width/depth), scale bar 500 µm). **C** Quantification of peak-to-peak amplitude and width of individual microwells upon 1 day of swelling in saline (****p<0.0001 by ANOVA and Bonferroni’s multiple comparisons test, n = 11 (Small, Med) and n = 10 (Large), see supplementary figure 4 for quantifications up to day 14). **D** Simulation of iAT2 cell localization upon seeding (cross section of center of microwell) and representative images of iAT2s (∼2 cells per µm^2^) seeded into microwells of different sizes at 1 hour (cell scale 12.5 µm (simulation) and scale bar 1 mm (microwells). **E** Quantification of projected alveolosphere areas at 14 days in culture (****p<0.0001 by ANOVA and Bonferroni’s multiple comparisons test, n = 27 (Small), 47 (Med), 41 (Large)). **F** Quantification of percentage of live area (quantified by Calcein AM (live cells) and Ethidium homodimer (dead cells) staining) at 14 days in culture (****p<0.0001, ***p<0.001, ns = not significantly different by ANOVA and Bonferroni’s multiple comparisons test, averaged from 4 independent experiments).

To test whether the hydrogel microwells support alveolosphere formation, we seeded iAT2s with an average density of 2 cells/µm^2^ hydrogel surface area in alveolosphere media. In silico modeling predicted cell sedimentation into individual microwells and brightfield imaging confirmed the formation of differently sized and irregularly shaped aggregates within 1 hour upon seeding (Fig. 2D). Formation of alveolospheres was controlled by the microwell size and initial seeding density, as measured by the projected alveolosphere area and viability. Within 14 days, iAT2 aggregates gave rise to alveolospheres across all microwell hydrogels (Supplementary Fig. 6) and alveolosphere area depended on microwell size (Fig. 2E). Although the efficiency and area increased in large microwells, alveolospheres also showed significantly reduced cell viability (∼70%, Fig. 2F), with dead cells mostly located in the core of individual alveolospheres (Supplementary Fig. 7), suggesting that insufficient diffusion and nutrient supply limit iAT2 survival. These results indicate that the medium-size microwell culture, in the absence of Matrigel or laminin/entactin enrichment, enables viable alveolosphere formation with high efficiency.

### iAT2 seeding density controls alveolosphere growth

Having shown that alveolospheres form in microwells, we used medium-size patterns to assess how iAT2 seeding density regulates growth and AT2 progeny within these engineered environments. Recent studies in intestinal organoids suggest a minimal number of cells is required to generate epithelial structures^21^. In addition, the initial aggregate size may induce paracrine signaling that directs self-assembly and growth of alveolospheres^21^. Thus, we seeded iAT2s at an average density of 15, 75 and 750 cells per individual microwell and monitored alveolospheres growth. At day 14, alveolospheres formed through all conditions with polarized apical surfaces (Fig. 3A), which is consistent with the epithelial structures observed in 3D hydrogels (Fig. 1). During the process of iAT2 self-assembly, alveolospheres grew in size as indicated by the increasing alveolosphere areas with culture with starting populations of 75 cells per well and higher, whereas lower seeding densities failed to generate growing alveolospheres (i.e., 75 cells, Fig. 3B). Similar growth patterns were observed in smaller microwells, further supporting that alveolosphere assembly and growth depends on initial iAT2 seeding densities (Supplementary Fig. 8).

**Figure 3.**
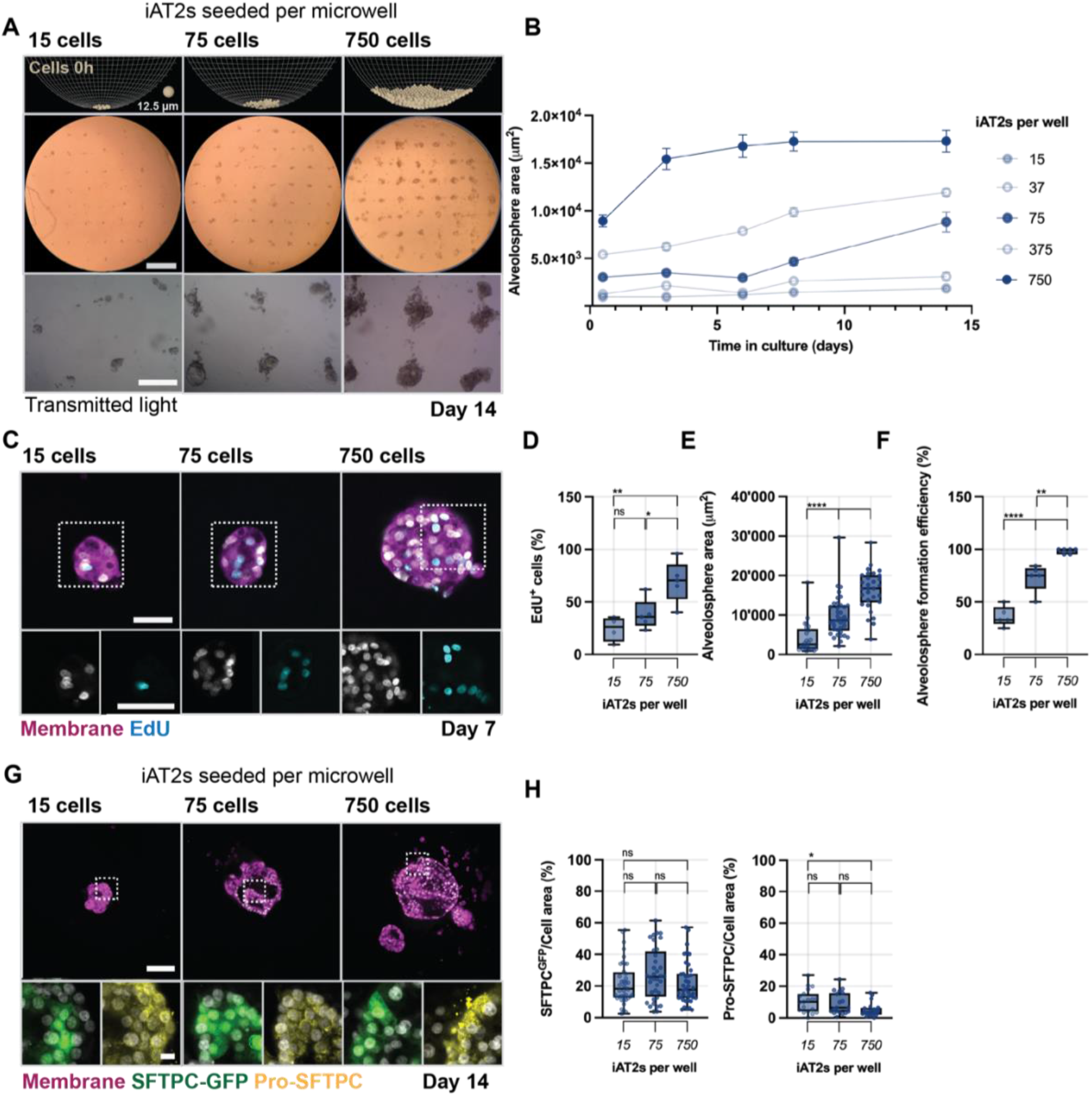
Effect of iAT2 seeding density on alveolosphere growth and fate. **A** Simulation of iAT2 cell localization upon seeding (cross section of center of microwell) and representative images of alveolospheres formed from 15, 75 and 750 iAT2s per individual medium size microwell (500 µm/200µm width/depth) at 14 days of culture (scale bars 1 mm (top) and 500 µm (bottom)). **B** Time course quantification of the projected area of alveolospheres formed from 15, 37, 75, 375 and 750 iAT2s over 14 days averaged from 3 independent experiments, see supplementary figure 6 for small size microwells). **C** Representative images of the incorporation of 5-ethynyl-2’-deoxyuridine (EDU) into alveolospheres over 7 days (cell mask membrane stain (magenta) nuclei (grey) and EdU (cyan), cell scale 12.5 µm (simulation), scale bars 100 µm (inset 10 µm)). **D** Quantification of the percentage of EdU^+^ nuclei at 7 days in culture, averaged from 4 independent experiments (*p<0.05, **p<0.01, ns = not significantly different by ANOVA and Bonferroni’s multiple comparisons test). **E** Quantification of the area of alveolospheres at 14 days in culture (****p<0.0001 by ANOVA and Bonferroni’s multiple comparisons test, n = 30 (15 iAT2s), n = 39 (75 iAT2s), and n = 31 (750 iAT2s). **F** Quantification of the percentage of microwells containing alveolospheres at 14 days of culture, averaged from 4 independent experiments (**p<0.01, ***p<0.001 by ANOVA and Bonferroni’s multiple comparisons test). **G** Representative images of SFTPC^GFP+^ and Pro-SFTPC expression in alveolospheres at 14 days of culture (cell mask membrane stain (magenta) nuclei (grey), SFTPC^GFP+^ (green), and pro-SFTPC (yellow), scale bars 100 µm (inset 10 µm). **H** Quantification of SFTPC^GFP+^ and pro-SFTPC expression per alveolosphere area (cell membrane stain) at 14 days (*p<0.05, ns = not significantly different by ANOVA and Bonferroni’s multiple comparisons test, n = 40 alveolospheres (SFTPC^GFP+^), n = 20 alveolospheres (pro-SFTPC).

The ability of iAT2s to generate alveolospheres is driven by their self-renewal potential and capacity to proliferate^9^, which may be influenced by the cell seeding density. EdU incorporation was used to visualize proliferating cells during the initial 7 days of culture (Fig. 3C). While we observed proliferating iAT2s throughout all conditions, the percentage of EdU+ cells was increased for alveolospheres formed from higher cell seeding densities (Fig. 3D). The increase in proliferative capacity further resulted in larger diameter alveolospheres at day 7 (Fig. 3E), suggesting the influence of cell-cell contact and paracrine signaling. In addition, quantification of alveolosphere forming efficiency revealed that individual microwells retained 36±14% of the alveolospheres with low cell seeding densities (15 cells), whereas greater efficiencies of 73±21-98±3% were maintained in microwells at higher seeding densities of 75 and 750 cells (Fig. 3F).

We next sought to determine whether the microwell culture method supported AT2 progeny of alveolospheres, focusing on the fluorescent reporter and expression of surfactant protein C (pro-SFTPC), which is highly specific to AT2 cells^26^. At day 14, we observed SFTPC^GFP+^ expressing cells in all alveolospheres that were also expressing pro-SFTPC, as identified by immunostaining (Fig. 3G). Varying the initial cell seeding density had little influence on SFTPC^GFP+^ levels, which ranged from 20-30%; however, lower cell seeding densities increased pro-SFTPC expression (Fig. 3H). This is consistent with a previous study that altered iAT2 plating densities to enhance AT2 progeny within Matrigel^25^.

### Microwell hydrogels maintain alveolosphere function

We next sought to define the cellular heterogeneity of alveolospheres and their AT2 progeny using single-cell RNA sequencing (scRNA-seq) and protein identification. To assess the functionality of microwell-cultured alveolospheres we used one condition (75 cells) in comparison to Matrigel cultured alveolospheres. At 14 days, cells clustered into 7 different clusters as visualized by Uniform Manifold Approximation and Projection (UMAP) and we identified several AT2 specific markers expressed by cells cultured under both conditions, including SFTPC and SFTPB (Fig. 4A, Supplementary Fig. 9). This data suggests that cells within alveolospheres similarly represent AT2-like cells. Interestingly, a greater proportion of the total cells in microwell conditions maintained expression of mature AT2 markers, suggesting microwell hydrogels may provide some advantages over Matrigel in terms of maintaining AT2 fate in alveolospheres.

**Figure 4.**
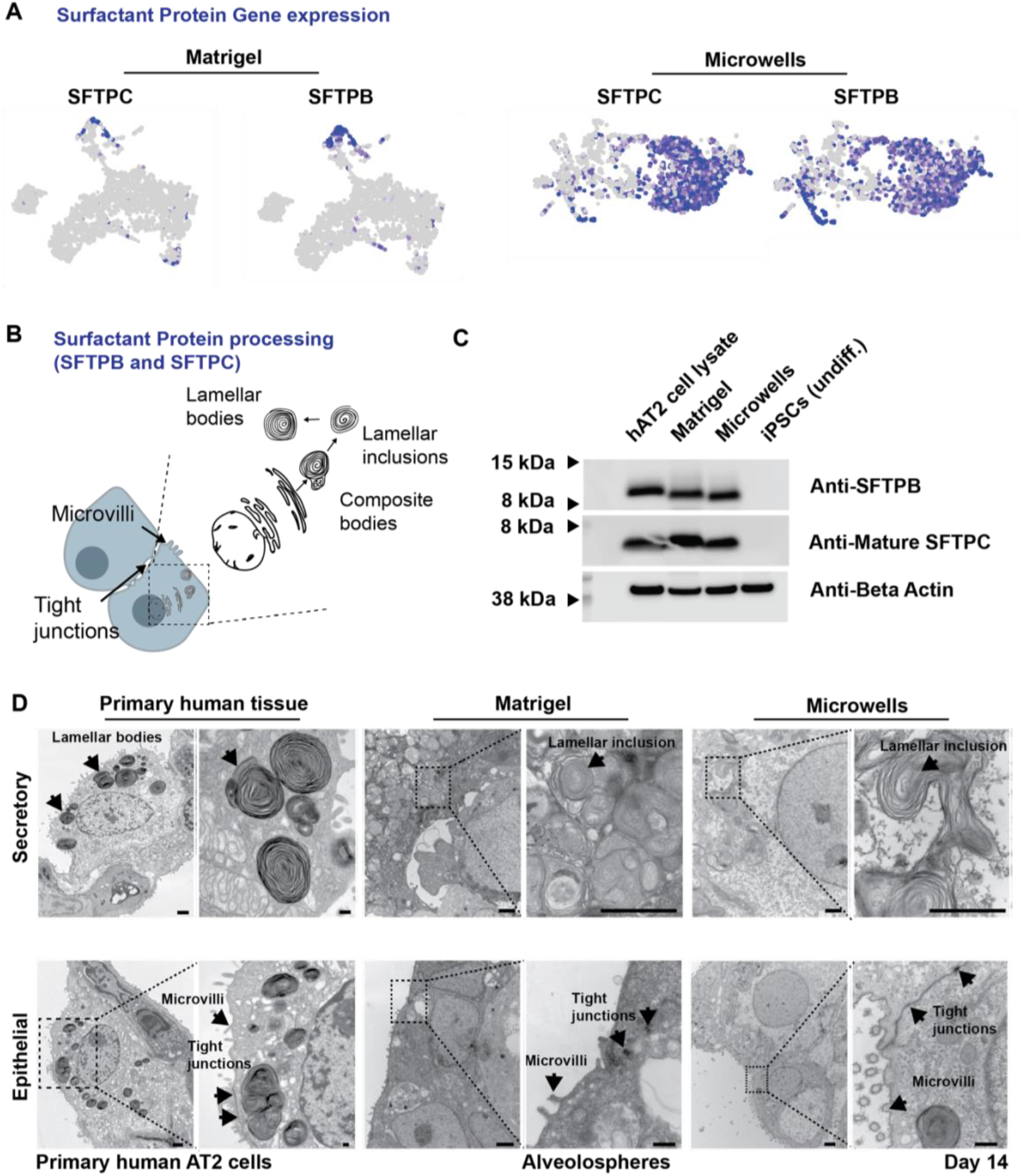
Characterization of microwell cultured alveolospheres. **A** UMAP representation of the expression of the AT2 markers surfactant protein C (SFTPC) and B (SFTPB) within alveolospheres at day 14 cultured in Matrigel and atop microwells. **B** Schematic illustrating the ultrastructural characteristics of differentiated AT2 cells, including epithelial cell differentiation (cell contacts with neighboring cells through tight junctions at the apical part of the lateral cell membrane and microvilli on the apical cell membrane) and secretory differentiation (Surfactant Protein B (SFTPB) and Surfactant Protein C (SFTPC) storage organelles and lamellar bodies that are mature or still in the process of maturation, defined as lamellar inclusions). **C** Western Blot for SFTPB, mature SFTPC and internal loading control Beta-Actin on human AT2 lysate, alveolospheres at day 14 cultured in Matrigel and atop microwells, as well as undifferentiated iPSCs. **D** Representative transmission electron microscopy of AT2 cells from primary human lung tissue samples and alveolospheres at day 14 cultured in Matrigel and atop microwells (scale bars 1 µm, inset 200 nm).

In addition, previous studies showed that iAT2 cells in Matrigel exhibit some aspects of AT2 cell maturation including apical tight junctions, apical microvilli, and the expression of lamellar-like bodies^9^ (Fig. 4B). Therefore, we next assessed whether cells were able to process pro-SFTPs to mature SFTPB and SFTPC proteins that are exclusive to AT2 cells and essential components of surfactant^26,27^. As a comparison, proteins were also extracted from human primary AT2 cells, alveolospheres cultured in Matrigel, and undifferentiated iPSCs. Western blots immunostained with antibodies that recognize the fully mature 8-kDa form of SFTPB and SFTPC revealed production of mature forms of each protein. Primary human AT2 controls and cells from both Matrigel and microwell-cultured alveolospheres also expressed the mature 8-kDa SFTPB and SFTPC proteins, whereas no staining of undifferentiated iPSCs was detected (Fig. 4C). As such, the microwell-cultured alveolospheres seem to efficiently process surfactant proteins into their mature form.

Next, we assessed whether the iAT2s in alveolospheres also formed lamellar bodies, the functional organelles in which surfactant is stored before exocytosis into the air spaces to form a phospholipid-rich film at the air-liquid interface^27^. Using transmission electron microcopy (TEM) analysis of primary human AT2 cells, these organelles are characterized by a tight packing of lipid lamellae into lamellar bodies. Similarly, Matrigel and microwell-cultured alveolospheres revealed lamellar-body like inclusions with a subset of inclusions expressing dense cores indicating the ongoing process of maturation (Fig. 4D). Importantly, microwell-cultured alveolospheres further showed typical characteristics of epithelial differentiation, including the formation of tight junctions at the apical part of the lateral cell membrane and microvilli on the apical cell membrane (Fig. 4D). These results suggest that alveolospheres when cultured in microwell hydrogels contain cells that form functional lamellar body-like inclusions, consistent with Matrigel cultures and findings reported in mature AT2 cells *in vivo*^26,28,29^.

### Orthotopic transplantation of human alveolar progenitors into murine lungs

Having shown that microwell-cultured alveolospheres retain adult AT2 phenotypic markers *in vitro*, we next sought to assess their viability and differentiation capacity *in vivo*. Recent studies have shown orthotopic transplantation of lung epithelial cells into injured lungs as a functional tool to interrogate the responsiveness of cells to *in vivo* signaling cues^30–34^. Here, we used bleomycin induced tissue injury, characterized by spatial AT2 cell loss within damaged regions^35^ (Supplementary Fig. 10). Immunodeficient non-obese diabetic (NOD) severe combined immunodeficiency (SCID) gamma (NSG) mice were injured with bleomycin at day 0, followed at day 10 by intranasal inhalation of alveolar progenitors derived from microwell and Matrigel-derived alveolospheres upon digestion (Fig. 5A). Immunostaining of human-specific nuclear and mitochondrial markers at day 14 post-transplantation showed discrete clusters of human cells, predominantly at the border of damaged alveolar regions in each of the recipient mice (n = 3 per group). Human cell clusters appeared to adopt similarity to neighboring alveoli and retain at least some alveolar differentiation, as indicated by some cells being positive for pro-SFTPC (Fig. 5B). Furthermore, staining for Ki67 expression revealed several proliferating cells that were located across the human cell clusters ((Fig. 5C). When analyzing cell clusters and percentage of Ki67+, we observed minimal differences between cells transplanted from alveolospheres cultured in Matrigel and microwells (Supplementary Fig. 11). The lack of Keratin 5 expression in both conditions further confirmed that transplanted cells did not trans-differentiate into basal cells (Supplementary Fig. 12). Although in contrast to recent findings on the differentiation of transplanted primary AT2 cells into basal cells^32^, the ability of cells to respond to the *in vivo* niche may depend on their origin (e.g., primary vs iPSC-derived), further highlighting the potential of the orthotopic transplantation assay to probe cell plasticity. We did not detect RAGE (receptor for advanced glycation endproducts), suggesting minimal differentiation into type 1 alveolar epithelial cells. Explanations for the lack of differentiation could be that the *in vitro* culture conditions are not supportive of AT1 differentiation,^9^ the early time points upon transplantation, or the incompatibility between murine and human growth factors / ECM components that are required for efficient AT1 fate adoption. These findings suggest that alveolospheres when cultured within engineered microwell hydrogels can retain their proliferative capacity upon transplantation into injured murine lungs, consistent with results in Matrigel and previous observations^36^. However, the lack of AT1 cells in both culture conditions as well as transcriptomic differences when compared to primary AT2 cells^9^ indicate that further optimization is still required for future applications, including cell-based therapies and disease modeling studies.

**Figure 5.**
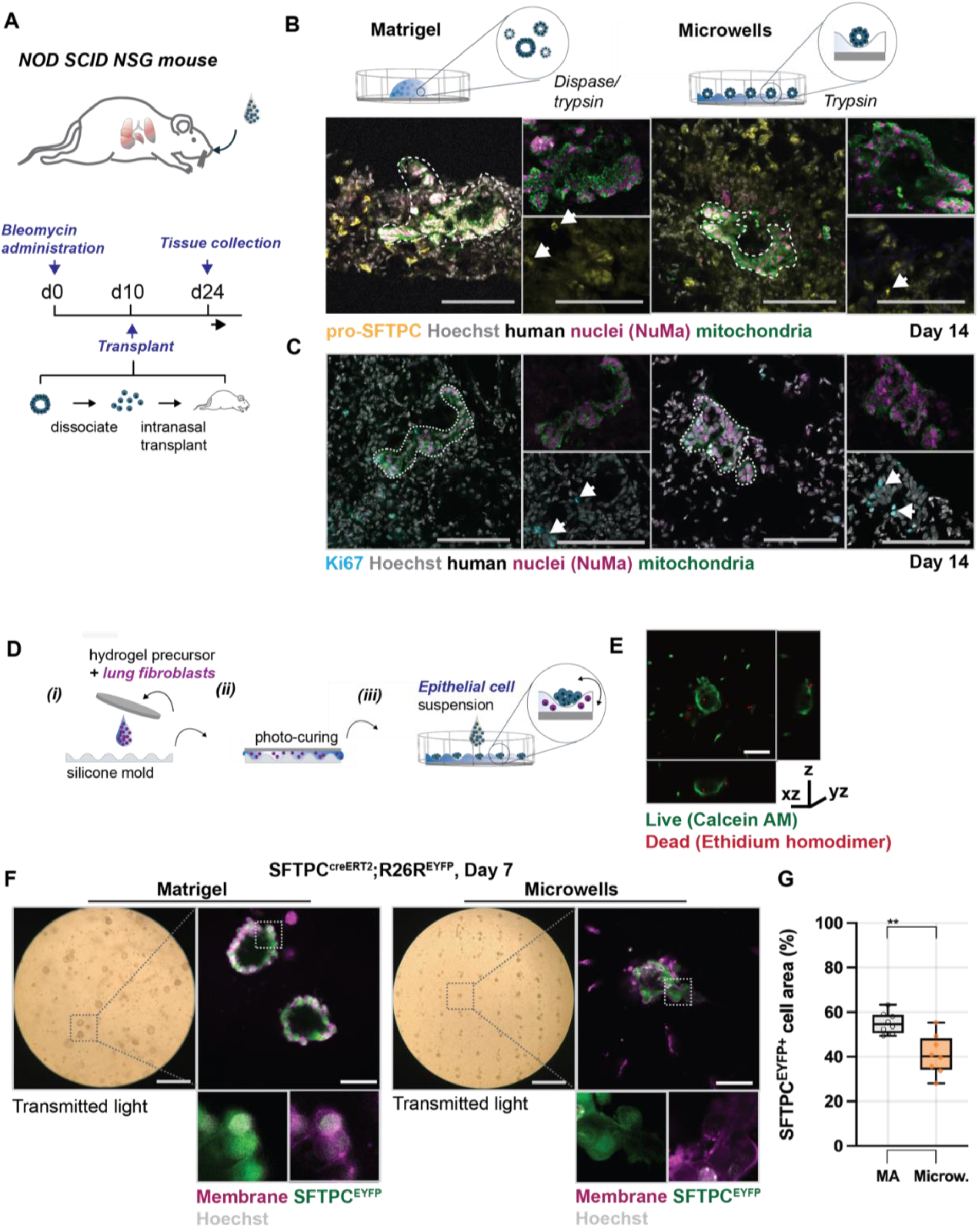
Applications of microwell cultured alveolospheres. **A** Schematic and transplant timeline of orthotopic transplantation of dissociated alveolospheres into bleomycin-injured, immunodeficient mouse lungs. **B** Representative images of the survival of cells derived from alveolospheres upon culture in Matrigel or atop microwells (nuclei (grey), pro-SFTPC (yellow), human nuclear marker (magenta), and human mitochondrial marker (green), scale bars 100 µm). See supplementary Fig. 10 for quantification of the transplanted cell area. **C** Representative images of the engraftment of cells derived from alveolospheres upon culture in Matrigel or atop microwells (nuclei (grey), Ki67 (cyan), human nuclear marker (magenta), and human mitochondrial marker (green), scale bars 100 µm). See supplementary Fig. 10 for quantification of percentage of Ki67^+^ cells per transplanted cell area. **D** Schematic illustrating the design of microwell hydrogels with encapsulated mouse fibroblasts and epithelial cells seeded atop. (i) Norbornene-modified hyaluronic acid hydrogel precursor solution was mixed with mouse fibroblasts (5×10^6^/mL) atop of a silicone molds and (ii) crosslinked through ultra-violet light-initiated thiol-ene reaction with a thiolated coverslip placed on top, flipped and (iii) followed by seeding epithelial cells atop. **E** Representative image of Calcein AM (live cells) and Ethidium homodimer (dead cells) staining of mouse alveolospheres formed in microwells at 7 days (medium: 500 µm/200 µm (width/depth)), scale bar 100 µm. **F** Representative images of SFTPC^CreERT2^; R26R^eYFP^ organoids formed in Matrigel (mixed with murine fibroblasts in a 1 to 10 ratio) and atop of hyaluronic acid hydrogel microwells (5% Matrigel, 5×10^6^ fibroblasts/mL, 21 kPa) with a seeding density of 75 SFTPC^CreERT2^; R26R^eYFP^ cells at 7 days (cell mask membrane stain (magenta) nuclei (grey) and SFTPC^EYFP+^ expression (green), scale bars 1 mm (transmitted light), 100 µm (SFTPC^GFP^)). **G** Quantification of SFTPC^EYFP+^ expression per cell area (cell membrane stain) of mouse AT2 organoids formed in Matrigel and atop of microwells at 7 days averaged from 3 independent experiments, **p<0.01 by unpaired two-tailed *t*-test).

### Microwell hydrogels enable culture of primary alveolospheres

While iPSC-derived alveolospheres have evolved as a reliable source of human AT2 cells, generation of primary alveolospheres continues to be critical, including for studies on mesenchymal to epithelial crosstalk and AT2 to AT1 differentiation^3,7,37^, which are currently not addressable in human alveolospheres^9^. Similarly, existing primary alveolosphere culture models depend almost exclusively on Matrigel as a 3D matrix, limiting their applicability for many downstream applications^12,38^. Thus, to further validate our microwell hydrogel system, we next adapted our design for the generation and culture of primary murine alveolospheres. Commonly, co-culture with mesenchymal cells is necessary as stromal support for alveolosphere development^7^. To address this, hydrogel microwells were fabricated with murine lung fibroblasts that are embedded during crosslinking, enabling mesenchymal to epithelial cell crosstalk during culture (Fig. 5C). Fibroblast cells were encapsulated at a density of 5 x10^6^ mL^−1^ and we confirmed that embedded cells did not alter microwell fidelity (Supplementary Fig. 13). To generate alveolospheres, SFTPC^EYFP+^ AT2 cells were isolated from Sftpc^CreERT2^ R26R^EYFP^ adult mice (Supplementary Fig. 14) and seeded directly into fibroblast-laden hydrogel microwells (75 cells per well), following our established protocol. Within 7 days, isolated AT2 cells assembled into lumenized structures that were physically separated from the embedded fibroblasts while maintaining high cell viability (Fig. 5D). Alveolospheres formed in the majority of individual microwells (∼75%, Fig. 5E), consistent with the efficiency of iAT2-derived alveolospheres (Fig. 3F,G); however, Sftpc^EYFP+^ expression levels were slightly lower when compared to Matrigel controls (Fig. 5F). We also observed that direct co-culture through the mixing of fibroblasts and SFTPC^EYFP+^ AT2s resulted in similar alveolospheres, whereas no alveolospheres were found in the absence of fibroblasts (Supplementary Fig. 15), supporting the importance of mesenchymal signaling during the formation of primary alveolospheres. These findings suggest that microwell hydrogels can be used for the generation of primary alveolospheres in a Matrigel-free environment, critical for a range of applications such as in modeling disease and probing mesenchymal to epithelial cell crosstalk.

### Outlook

We have designed a versatile Matrigel-free culture system to generate lung alveolospheres either when embedded within 3D hydrogels or seeded atop microwell hydrogels. Previous studies to engineer organoids within microwell hydrogels have focused on the development of intestinal, pancreatic and cancer organoids^21–23^, but there has been little work to use engineered culture systems for lung organoids. Often, customized engineering technologies or hydrogel chemistries are necessary to fabricate microwell hydrogels, which prevents their translation across groups. By using commercially available culture dishes with pre-formed microwells, we fabricated cytocompatible hydrogels with evenly spaced microcavities, which enabled the generation and culture of functional alveolospheres. Our data indicate that within microstructured hydrogels, human alveolospheres maintain their proliferative and differentiation capacity. This approach further supports the versatility of embedding other cell types, such as lung fibroblasts, into the hydrogel to enable the formation of primary mouse alveolospheres. The supportive mesenchyme was found to provide paracrine signals that are essential for alveolar epithelial cell self-assembly and organization. Thus, the functional encapsulation of fibroblast populations in engineered, functional alveolosphere cultures provides new perspectives on disease modeling. We anticipate that by modulating specific culture conditions such as fibroblast lineages, hydrogel stiffness and composition, the microwell culture system can be used to study the role of mesenchymal-to-epithelial cell communication^37^ and biophysical signaling during *in vitro* alveologenesis.

AT2 functionality was tested through analysis of surfactant protein processing and survival upon orthotopic transplantation of dissociated alveolospheres into lungs of immunocompromised mice. When microwell-cultured alveolospheres were transplanted into injured murine lungs, dissociated cells adopted alveoli-like structures while maintaining their proliferative capacity, and this was similar to alveolospheres formed within Matrigel. Previous work has reported data on the capacity of orthotopic transplanted epithelial progenitor cells to survive in injured murine lungs^31,32,34^. However, the technology described here could hold promise as a means to expand functional epithelial cells in Matrigel-free conditions for therapeutic applications. Further studies are required to probe whether cells are structurally and functionally integrated into mouse lungs, such as clonality and long-term replacement of injured epithelium; yet, to date, little is known regarding methods that confirm functional engraftment, or even the definition of engraftment in this context^33^. Although orthotopic transplantation assays are not yet sufficient to assess the contribution of transplanted cells to organ function, it provides a powerful platform to probe iPSC-derived lung epithelial cell viability, plasticity, and ability to incorporate into a compatible host tissue. Finally, although we observed lamellar inclusions and the functional capacity to process SFTPB and SFTPC to their mature form, extracted proteins from both conditions present relatively lower amounts when compared to primary AT2 cells. While these differences might, in part, be explained by the effect of *in vitro* culture and sample processing, issues of limited characterization remain to be eliminated, further highlighting the need for culture platforms that support culture of both iPSC-based and primary AT2 cells.

Taken together, there are currently no strategies for the defined culture of lung alveolospheres in Matrigel-free conditions. Thus, the microwell hydrogels described herein provide means as an accessible culture system for the generation and maintenance of primary and iPSC-derived lung progenitors, which may be extendable to other epithelial progenitor and stem cell aggregates.

## Methods

### Animals

The R26R^EYFP+^ mouse line was purchased from Jackson Laboratories and the Sftpc^CreERT2^ mouse line generously provided by Dr. Hal Chapman, and genotyping information has been previously described^39^. All mice were maintained on a mixed background and were 3-5 weeks of age for experiments in this study. NOD.CgPrkdc^scid^ Il2rg^tm1Wj^l/SzJ (NSG mice) were utilized as recipients for all transplantation experiments. Mice were 8-10-week-old mice with male and female in equal proportions. Experiments were not blinded to mouse age or sex. All experiments were carried out under the guidelines set by the University of Pennsylvania’s Institutional Animal Care and Use Committees and followed all NIH Office of Laboratory Animal Welfare regulations.

### Tamoxifen induction

For lineage tracing, tamoxifen (Sigma-Aldrich) was dissolved in corn oil at a concentration of 20 mg/ml. One-week prior sacrifice, mice were administered tamoxifen (200 mg/kg) by oral gavage.

### Bleomycin Lung injury

10 days prior cell transplantation, mice were first anesthetized using 3.5% isoflurane in 100% O_2_ via and anesthesia vaporizer system. Mice were intranasally administered 4 mg/kg body weight bleomycin sulfate in a total volume of 30 µL PBS. Only injured mice that lost ∼10% of their starting body weight by day 4 post injury and survived to the time of transplant were considered to be adequately injured and used for all transplantation experiments.

### Orthotopic cell transplantation

Dissociated alveolosphere were administered via intranasal inhalation to NSG mice as previously described^31^. Recipients received 1.2 million iAT2 cells. Recipient mice were anesthetized with 3.5% isoflurane in 100% O_2_ via an aesthesia vaporizer system and were intranasally administered cells by pipetting 30 µL single cell suspension in PBS (containing 1% penicillin-streptomycin) onto the nostrils of anesthetized mice visually confirmed by agonal breathing.

### Lung tissue harvest

Following sacrifice via isoflurane overdose, lungs were inflated at a constant pressure of 125 cm H2O with 3.2% paraformaldehyde (PFA) for 30 minutes followed by incubation in 3.2% PFA for another 30 minutes at room temperature. Fixed lungs were then washed in multiple PBS washed over the course of 1 hour at room temperature, followed by an overnight incubation in 30% sucrose shaking at 4°C, and then a 2-hour incubation in 15% sucrose shaking at 4°C, and then a 2 hour incubation in 15% sucrose 50% Tissue-Tek^®^ O.C.T. Compound at room temperature. Finally, fixed lungs were embedded in O.C.T. by flash freezing with dry ice and ethanol. 10 µm tissue sections were cut on a cryostat, followed by fixation in 4% PFA for 5 min, rinsed three times with PBS, and blocked in blocking solutions (PBS + 1% bovine serum albumin + 5% horse serum + 0.1% Triton X-100 for 45 minutes. Slides were incubated in primary antibodies in blocking solution overnight at 4C, washed three times with PBS + 0.1% Tween-20 and subsequently incubated with secondary antibodies for 2 hours at room temperature. Slides were then washed with PBS Tween-20 prior to incubation in 1 µM Hoechst. For Hematoxylin and eosin (H&E) staining, lungs were prepared as described above and paraffin embedded using standard procedures^40^.

### Lung cell isolation

One week post tamoxifen induction, mice were euthanized by CO2 inhalation. The chest cavity was exposed, and the right ventricle perfused with ice-cold PBS to clear lungs from blood. Lungs were removed and minced with a razor blade. Minced lungs were placed in a digestion solution containing 480 U/ml Collagenase Type I (Life Technologies) 50 U/ml Dispase (Collaborative Biosciences) and 0.33 U/ml DNase (Roche). Lungs were incubated in this solution for 45 min with frequent mixing in a 37° C water bath. The obtained solution was then filtered through a 100 µm cell strainer (BD Falcon), ACK lysis buffer added to remove blood cells and cell pellet resuspended in sterile PBS containing 1% FBS and 1 mM EDTA. Cells were stained for 30 min with Epcam-APC (1:1000), washed once prior Flow Cytometry sorting. Cell sorting was performed on a FACS Jazz (BD Biosciences) and sorted cells collected in sterile PBS containing 1% FBS and 1 mM EDTA. Lung fibroblasts were obtained as previously described. Briefly, minced lungs were digested as described above, passed through a 100 µm cell strainer, resuspended in ACK lysis buffer before passing through a 40 µm cell strainer. The resuspended cells were cultured on tissue culture plastic with DMEM-F12 plus 10%FBS. Media was refreshed after 1 hour and subsequently every other day. Lung fibroblasts were maintained for no more than three passages.

### iPSC cell line generation and maintenance

All experiments using human iPSC lines were approved by the Institutional Review Board of the University of Pennsylvania. The human pluripotent stem cell line RUES2 was obtained from the University of Pennsylvania iPSC Core Facility. To generate a stable human-derived alveolar epithelial type II-like cells (iAT2) cell line, a EGFP cassette was knocked-in on one allele of the SFTPC gene (RUES2-SFTPC-EGFP) and differentiation performed as previously described^25^. Upon differentiation, iAT2 cells were passaged every 14 days and sorted for GFP+ cells using a BD FACSJazz™ cell sorter. GFP+ cells were plated as single cells using 90% Matrigel and a density of 400 cells/µl. Cells were cultured in Iscove’s Modified Dulbecco’s Medium (IMDM)/Ham’s F12 media supplemented with 3 µg CHIR99021, 10 ng/mL KGF, 50 µM dexamethasone, 0.1 mM 3-Isobutyl-1-methylxanthine (IBMX), 43 µg/mL 8-Bromo-cAMP and primocin (CK+DCI medium), containing 10 µM Y-27632 for the first 48 h after plating, followed by 5 days in K+DCI (without CHIR99021) and 7 days in CK+DCI. Alveolosphere were passaged every 14 days by digesting the Matrigel with 2 mg/mL dispase for 1h at 37°C, followed by incubation in 0.25% Trypsin/EDTA for 10 min at 37°C to obtain a single cell suspension. Cell quantification and viability were assessed using Trypan blue. Finally, cells were mixed with Matrigel, 50 µL drops formed within 24 well plates and incubated for 30 min at 37°C and 5% CO_2_.

### Electron Microscopy

Alveolospheres were released from Matrigel using dispase or mechanically retrieved from microwells through several PBS washes, followed by fixation in 2.5% glutaraldehyde in 0.1% cacodylate buffer for at least 3 h at room temperature. Sample preparation was performed as recently reported^9^. Briefly, dehydration was performed with acetone on ice and graded ethanol series. Samples were then incubated in 100% acetone at RT for 2×10 min and in propylene oxide at RT for 2×15 min. Finally, samples were embedded in Embed-812 (Electron Microscopy Sciences), incubated in uranyl acetate and lead citrate and imaged with a JEOL 1010 electron microscope including a Hamamatsu digital camera (AMT Advantage image capture software).

### Western Blotting

Western blots were performed to detect processed SFTPC and SFTPB protein as previously described^41^. Briefly, total protein content of cell lysates was assayed by the Bradford method followed by SDS-PAGE and immunoblotting. Western blotting used a previously published polyclonal pro-SFTPB antiserum (“PT3-SP-B” at 1:3000 dilution), a commercially available mature SFTPC antibody (WRAB-76694; Seven Hills Bioreagents at 1:2500 dilutiion), and Beta-Actin (Sigma Aldrich A1978 at 1:10000 dilution) followed by an HRP-conjugated secondary antibody and visualization by enhanced chemiluminescence.

### Hydrogel preparation and seeding

#### Hydrogel synthesis

Norbornene-modified Hyaluronic acid (NorHA) was synthesized as described previously^42^. The degree of modification was 26% by ^1^H NMR (Supplementary Fig. 15). Enzymatically (metalloproteinase (MMP)) degradable di-thiolated peptides (GCNSVPMSMRGGSNCG) and thiolated cell-adhesive RGD peptides (GCGYGRGDSPG) were purchased from Genscript. NorHA hydrogels were fabricated by thiol-ene addition crosslinking with either ultraviolet (microwells) or visible light (3D hydrogels) and the photo-initiator lithium phenyl-2,4,6-trimethylbenzoylphosphinate (LAP, Colorado Photopolymer Solutions).

#### iAT2 encapsulation and culture

Hydrogel precursor solutions (4wt% polymer) were mixed with 1000 cells per µL (or as otherwise noted), 1 mM thiolated RGD and mixed with or without laminin/entactin (Corning, 354259) or Matrigel at different concentrations, and photo-polymerized with MMP-degradable peptide crosslinkers (400-500 nm, Omnicure S1500, Exfo) for 10 min at 10 mW cm^−2^. Gels were crosslinked as 50 µL droplets atop thiolated coverslips and cultured in 48-well plates^43^. Cells were cultured in CK+DCI medium with 10 µg/mL Y27 for the first 48 h.

#### Microwell fabrication and culture

Microwell replicate topographies were fabricated by moulding from cell culture surfaces (EZSPHERE^TM^) with different microwell width and depth. Briefly, poly(dimethylsiloxane) PDMS (Sylgard^TM^ 184, Ellsworth Adhesives, 10:1 ratio) was mixed with Hexanes (30% vol/vol), and polymerized for 2 h at 80°C. Hydrogel microwell topographies were fabricated through NorHA mixed with 1 mM thiolated RGD and crosslinked with MMP-degradable peptide crosslinkers (320-390 nm, Omnicure S1500, Exfo) for 5 min at 5 mW cm^−2^. For primary AT2 culture, mouse lung fibroblasts were encapsulated at 5 million cells/mL (or as otherwise noted). iPSC or primary mouse AT2 cells were added atop with different cell densities and cultured in CK+DCI medium (iAT2) or modified SAGM media as previously described^40^. Briefly, Small Airway Epithelial Cell Growth Basal Media (SABM, Lonza) was mixed with Insulin/Transferrin, Bovine Pituitary Extract, Gentamycin, and Retinoic Acid as well as 0.1 mg/mL Cholera Toxin (Millipore Sigma), 25ng/mL EGF (Peprotech), and 5% FBS. CK+DCI and SAGM media were supplemented with 10 µg/mL Y27 for the first 48 h.

### In silico modeling

To simulate initial cell seeding within microwells, Cinema 4D (C4D) rigid body dynamic simulations were used. Briefly, microwells were created according to various geometries and tagged as collider bodies, and cells (spheres, 12.5 µm) were tagged as rigid bodies and seeded into wells using simulated gravity. Cells were arrayed above the microwell with random seed points, and after settling, a Boole object was used to segment the simulation in half and rendering was carried out using C4D.

### Single cell RNA sequencing

Single cells suspensions from alveolospheres within Matrigel and microwells were prepared as outlined above. Cells were pelleted and counted by trypan blue, resuspended in sterile PBS containing 0.04% BSA for 10X Genomics, aiming for 10000 cells. Cells were loaded onto a GemCode instrument (10x Genomics, Pleasanton, CA, USA) to generate single-cell barcoded droplets (GEMs) according to the manufacture’s protocol. The resulting libraries were sequenced on an Illumina NovaSeq 6000 instrument.

### Analysis of scRNA-seq data

Reads were aligned and unique molecular identifier (UMI) counts obtained using STAR-Solo (v2.7.9a)^44^. Seurat (v4.0.1)^45^ was used for all downstream scRNA-seq analysis. Cells with less than 200 genes, greater than 2 Median absolute deviation above the median, and with potential stress signals of greater than 25% mitochondrial reads were removed. Data was normalized and scaled using the SCTransform function and adjusting for percent mitochondria, number of features per cell, and number of UMI per cell. Linear dimension reduction was done via PCA, and the number of PCA dimensions was evaluated and selected based on assessment of an ElbowPlot. The Uniform Manifold Projection (UMAP) data reduction algorithm was used to project the cells onto two dimensional coordinates. The Seurat function FeaturePlot was used create the UMAP gene expression plots.

### Statistical analysis and reproducibility

GraphPad Prism 9 software was used for statistical analyses. Statistical comparisons between two experimental groups were performed using two-tailed Student’s t-tests and comparisons among more groups were performed using one-way or two-way ANOVA with Bonferroni post hoc testing. All experiments were repeated as described in the text.

## Acknowledgements

This work was supported by funding from the NIH (K99-HL151670 to C.L., F32 DK117568 to M.D.D), and NSF (NSF STC program (CMMI): 15-48571 to C.L., M.D.D. and J.A.B.). M.O. acknowledges funding by the German Research Federation (DFG, SFB 1449 / B01). We would like to thank Florin Tuluc and staff at the Flow Cytometry Core Laboratory of Children’s Hospital of Philadelphia, Biao Zuo and his staff at the Electron Microscopy Core at the University of Pennsylvania, the iPSCs core at the University of Pennsylvania for the RUES2 cell line, Luis Rodriguez for fruitful discussions, and Gargi Palashikar for assistance with lung harvest.

## Author contributions

C.L., J.A.B., M.F.B., A.E.V, and E.E.M. conceived the ideas and designed the experiments. C.L. conducted the experiments on iPSC and primary alveolosphere culture and analysis, A.I.W. and A.E.V. on orthotopic transplantation, F.L.C. generated and maintained iPSC-derived AT2 cells, J.B.K. performed Western Blot analyses, K.S. obtained histology data on bleomycin-injured lungs, M.O. acquired EM data of human primary AT2 cells, M.C.B. and M.P.M performed scRNA-seq analysis, and M.D.D. obtained in-silico simulations. C.L., and J.A.B. wrote the manuscript, all authors read and edited the manuscript.

---

**Supplementary Figure 1:**
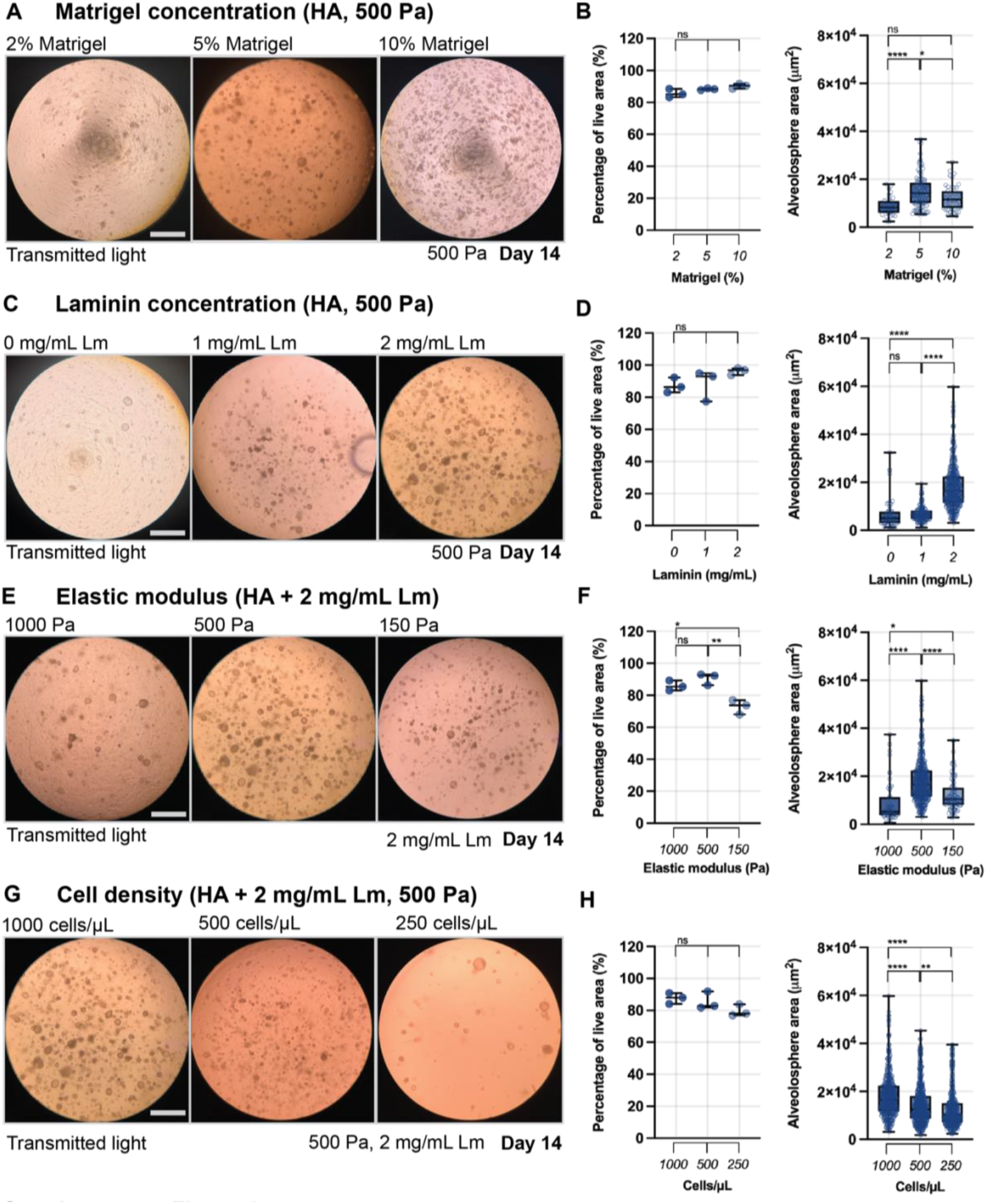
Influence of hyaluronic acid hydrogel composition, modulus and seeding density on viability and growth of alveolospheres. **A** Representative images of alveolospheres formed in hyaluronic acid hydrogels containing Matrigel (2%, 5% and 10% (vol/vol)) at 14 days (G’ 500 Pa with a seeding density of 1000 iAT2s/µL). **B** Quantification of percentage of live area (quantified by Calcein AM (live cells) and Ethidium homodimer (dead cells) staining) and the projected area of alveolospheres at 14 days in culture (****p<0.0001, *p<0.05, ns = not significantly different by ANOVA and Bonferroni’s multiple comparisons test, live area averaged from 3 independent experiments, n = 32 (2%), 106 (5%), 48 (10%)). **C** Representative images of alveolospheres formed in hyaluronic acid hydrogels containing laminin/entactin (0 mg/mL, 1 mg/mL, 2 mg/mL) at 14 days (G’ 500 Pa with a seeding density of 1000 iAT2s/µL). **D** Quantification of percentage of live area (quantified by Calcein AM (live cells) and Ethidium homodimer (dead cells) staining) and the projected area of alveolospheres at 14 days in culture (****p<0.0001, ns = not significantly different by ANOVA and Bonferroni’s multiple comparisons test, live area averaged from 3 independent experiments, n = 55 (0 mg/mL), 165 (1 mg/mL), 430 (2 mg/mL)). **E** Representative images of alveolospheres formed in hyaluronic acid hydrogels of different initial modulus (1000 Pa, 500 Pa, 150 Pa) at 14 days (2 mg/mL laminin/entactin with a seeding density of 1000 iAT2s/µL). **F** Quantification of percentage of live area (quantified by Calcein AM (live cells) and Ethidium homodimer (dead cells) staining) and the projected area of alveolospheres at 14 days in culture (****p<0.0001, **p<0.01, *p<0.05, ns = not significantly different by ANOVA and Bonferroni’s multiple comparisons test, live area averaged from 3 independent experiments, n = 53 (1000 Pa), 430 (500 Pa), 90 (150 Pa)). **G** Representative images of alveolospheres formed in hyaluronic acid hydrogels with different cell seeding densities (1000 cells/µL, 500 cells/µL, 250 cells/µL) at 14 days (G’ 500 Pa with 2 mg/mL laminin/entactin). **H** Quantification of percentage of live area (quantified by Calcein AM (live cells) and Ethidium homodimer (dead cells) staining) and the projected area of alveolospheres at 14 days in culture (****p<0.0001, **p<0.01, ns = not significantly different by ANOVA and Bonferroni’s multiple comparisons test, live area averaged from 3 independent experiments, n = 434 (1000 cells/µL), 630 (500 cells/µL), 410 (250 cells/µL)).

**Supplementary Figure 2:**
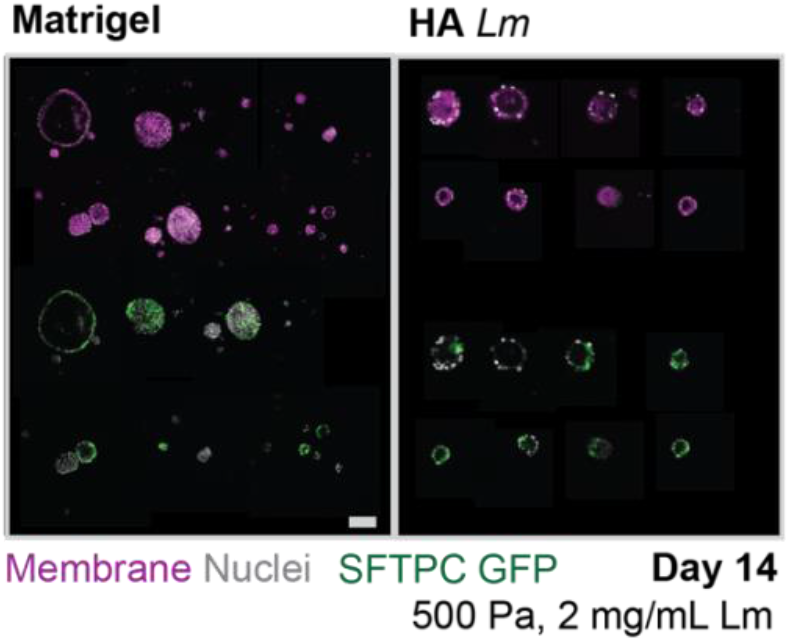
Expression of SFTPC^GFP+^ in alveolospheres formed in Matrigel and hyaluronic acid hydrogels with 2 mg/mL laminin/entactin. Representative images of SFTPC^GFP+^ alveolospheres formed in Matrigel and hyaluronic acid hydrogels with 2 mg/mL laminin/entactin (HA Lm, 500 Pa) with a seeding density of 1000 cells/µL at 14 days (cell mask membrane stain (magenta) nuclei (grey) and SFTPC^GFP+^ expression (green), scale bar 100 µm).

**Supplementary Figure 3:**
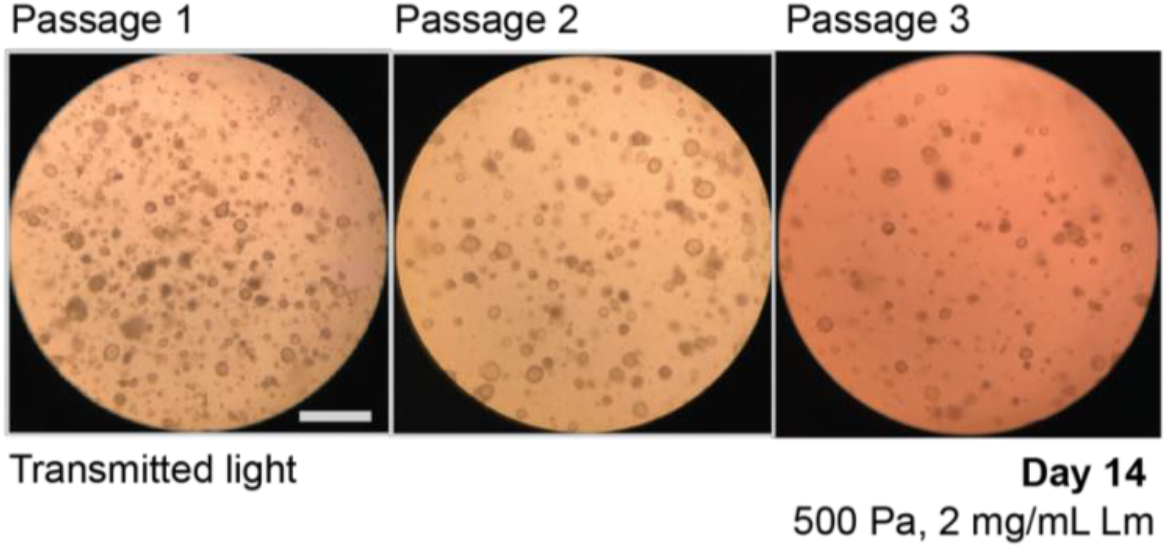
Passaging of al alveolospheres in hyaluronic acid hydrogels with 2 mg/mL laminin/entactin. Representative images of alveolospheres after serial passages in hyaluronic acid hydrogels with 2 mg/mL laminin/entactin (HA Lm, 500 Pa, 1000 cells/µL).

**Supplementary Figure 4:**
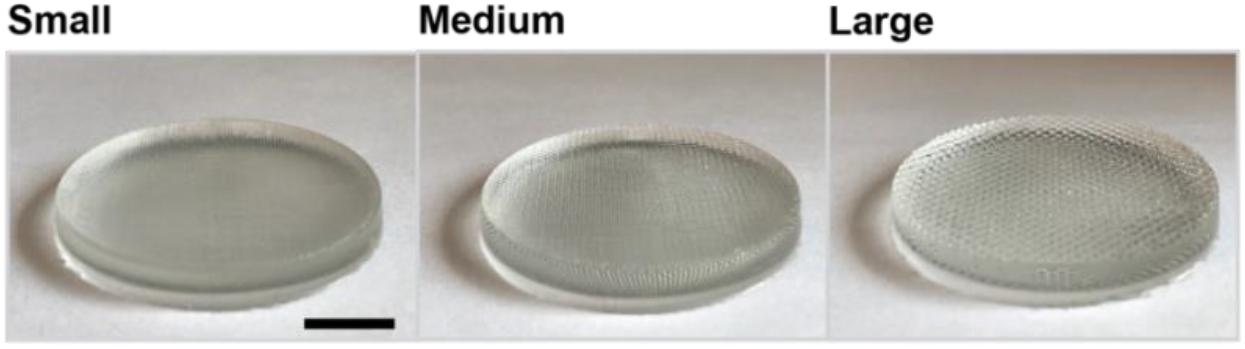
Silicone replica molds made from EZSPHERES^TM^ dishes. Representative images of silicone replica molds of different size that were fabricated from commercially available cell culture surfaces (EZSPHERES^TM^, small: 200 µm/100 µm, medium: 500 µm/200 µm, large: 800 µm/300 µm (width/depth), scale bar 1 mm).

**Supplementary Figure 5:**
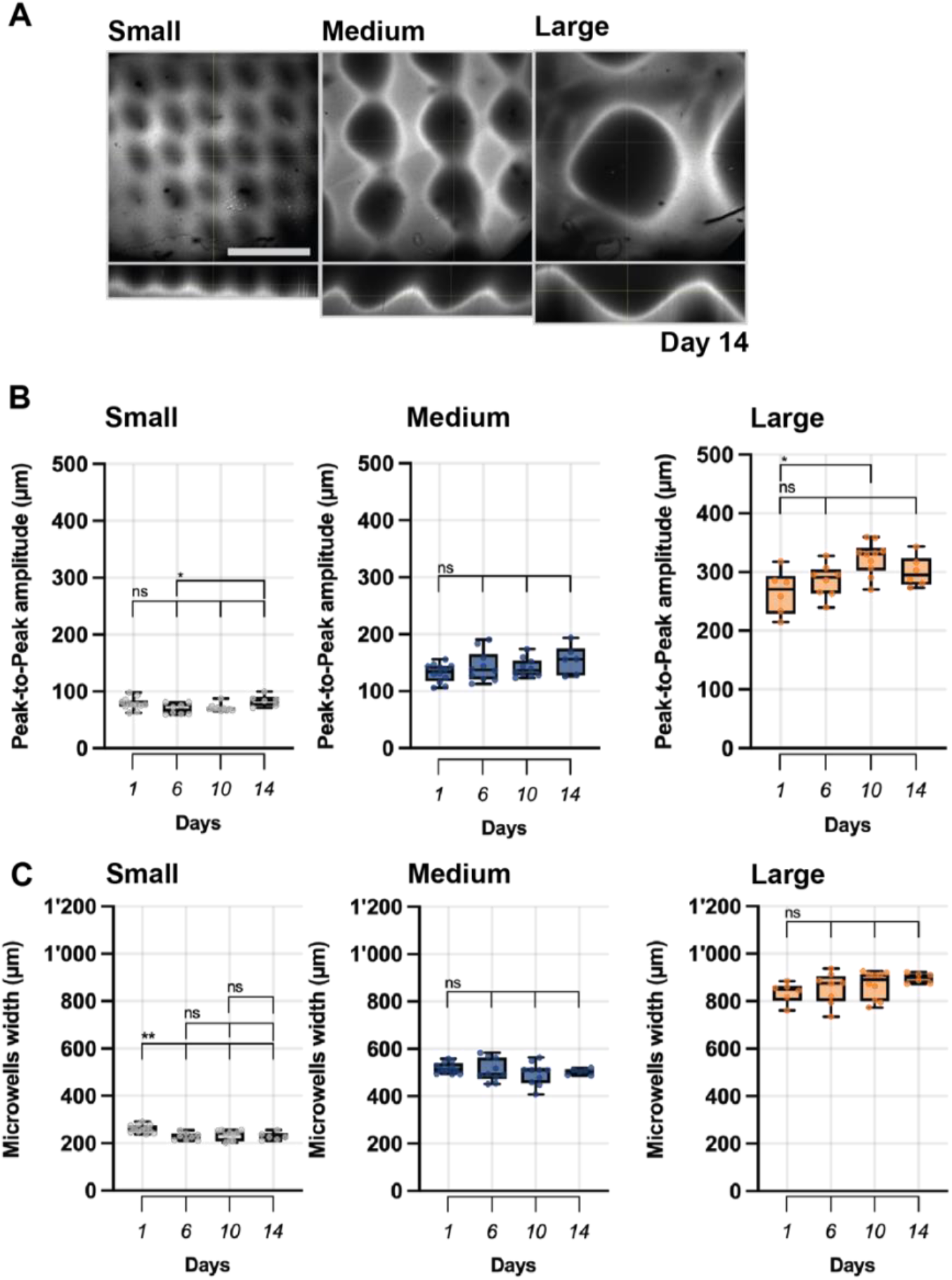
Microwell hydrogel stability and fidelity. **A** Representative images of microwell hyaluronic acid hydrogels modified with fluoresceine and fabricated from silicone molds with different width and depth upon 14 days of swelling in saline (small: 200 µm/100 µm, medium: 500 µm/200 µm, large: 800 µm/300 µm (width/depth), scale bar 500 µm). **B** Quantification of peak-to-peak amplitude of individual microwells during swelling in saline (day 1, 6, 10, 14), *p<0.05, ns = not significantly different by ANOVA and Bonferroni’s multiple comparisons test, n = 11 (Small, Med) and n = 10 (Larg). **C** Quantification of the width of individual microwells during swelling in saline (day 1, 6, 10, 14), **p<0.01, ns = not significantly different by ANOVA and Bonferroni’s multiple comparisons test, n = 11 (Small, Med) and n = 10 (Larg).

**Supplementary Figure 6:**
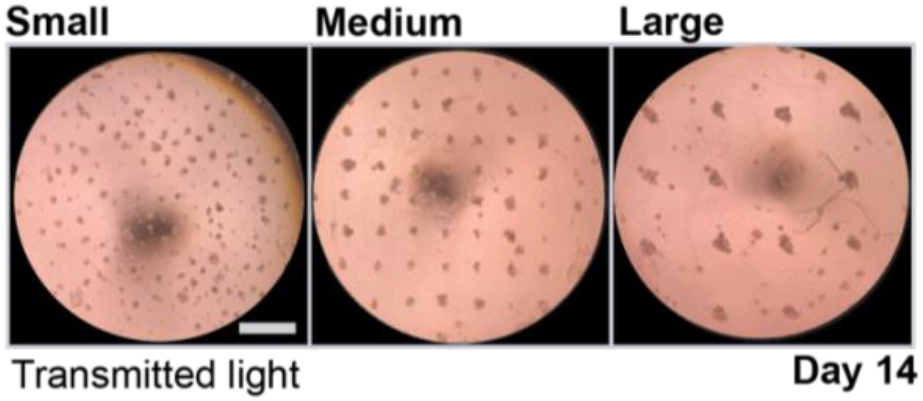
Formation of alveolospheres in microwells of different size. Representative images of alveolospheres formed in microwells of different size at 14 days (small: 200 µm/100 µm, medium: 500 µm/200 µm, large: 800 µm/300 µm (width/depth), scale bar 1 mm.

**Supplementary Figure 7:**
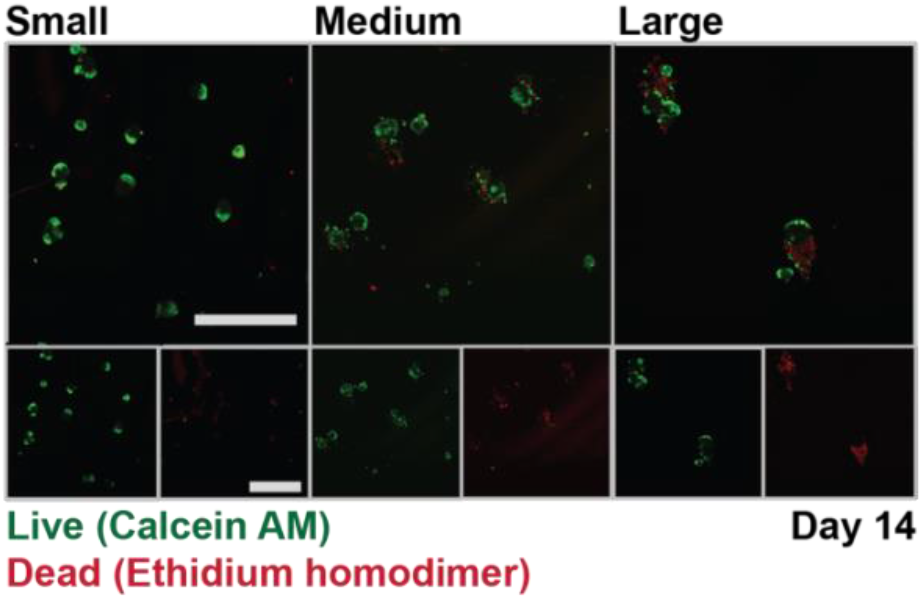
Viability of alveolospheres within microwells of different size. Representative images of Calcein AM (live cells) and Ethidium homodimer (dead cells) staining of alveolospheres formed in microwells of different size at 14 days (small: 200 µm/100 µm, medium: 500 µm/200 µm, large: 800 µm/300 µm (width/depth), scale bars 500 µm.

**Supplementary Figure 8:**
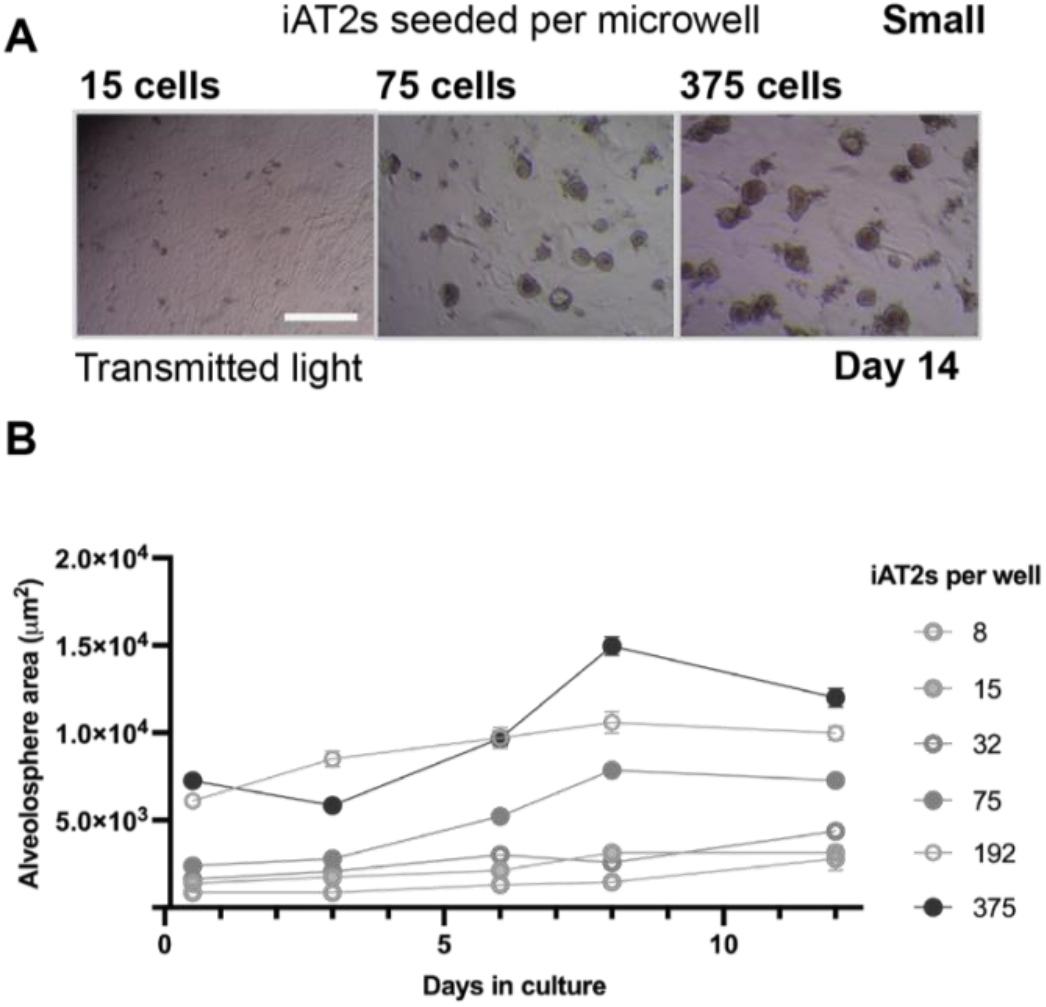
Formation of alveolospheres in small microwells. **A** Representative images of alveolospheres formed from 15, 75 and 375 iAT2s per individual medium size microwell (200 µm/100µm width/depth) at 14 days of culture (scale bar 500 µm). **B** Time course quantification of the projected area of alveolospheres formed from 8, 15, 32, 75, 192and 375 iAT2s over 14 days averaged from 3 independent experiments.

**Supplementary Figure 9:**
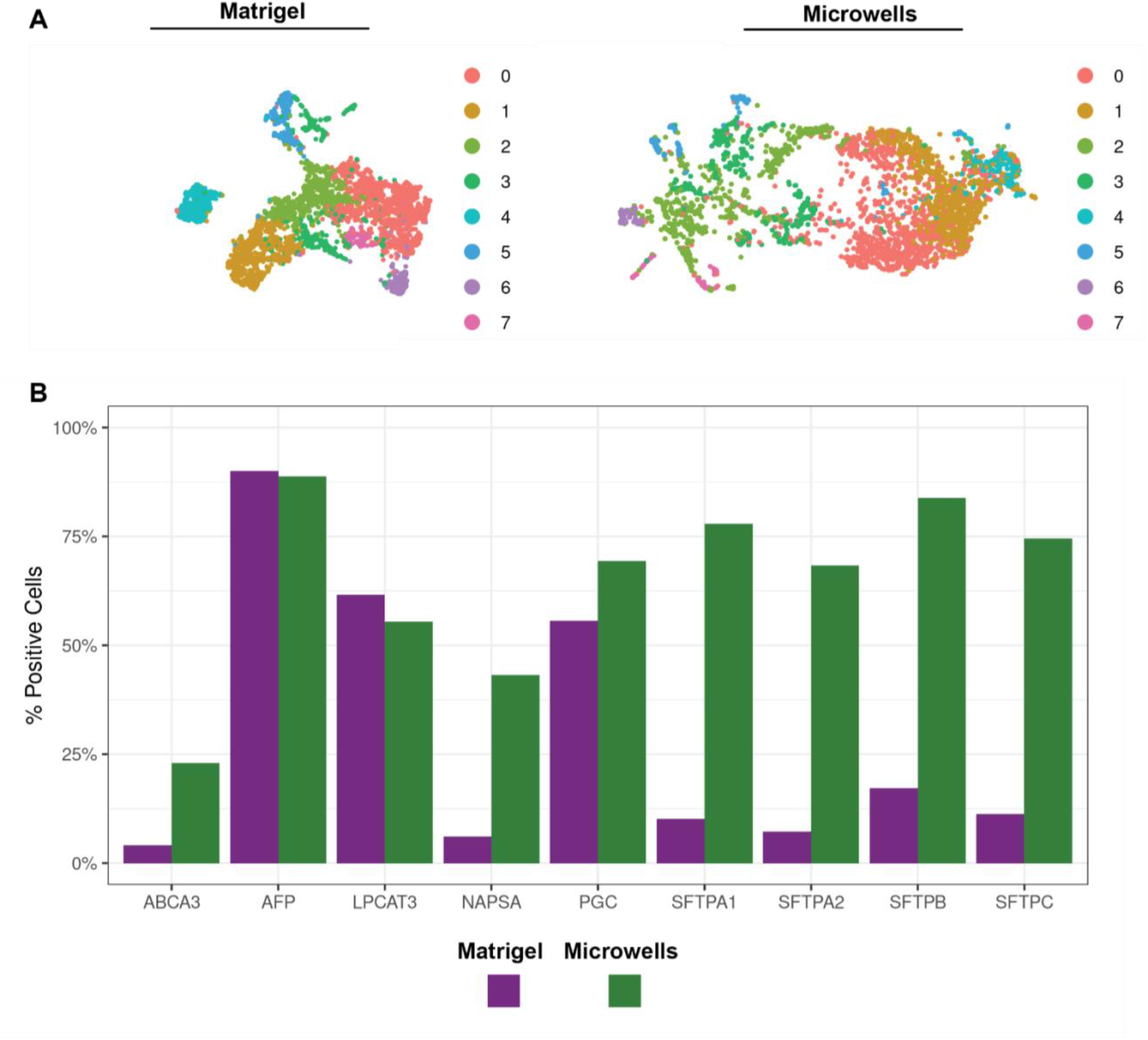
ScRNA-seq characterization of alveolospheres. **A** UMAP representation of 7 clusters and **B** gene expression proportion of AT-2 specific markers of cells within alveolospheres at 14 days cultured in Matrigel and atop microwells.

**Supplementary Figure 10:**
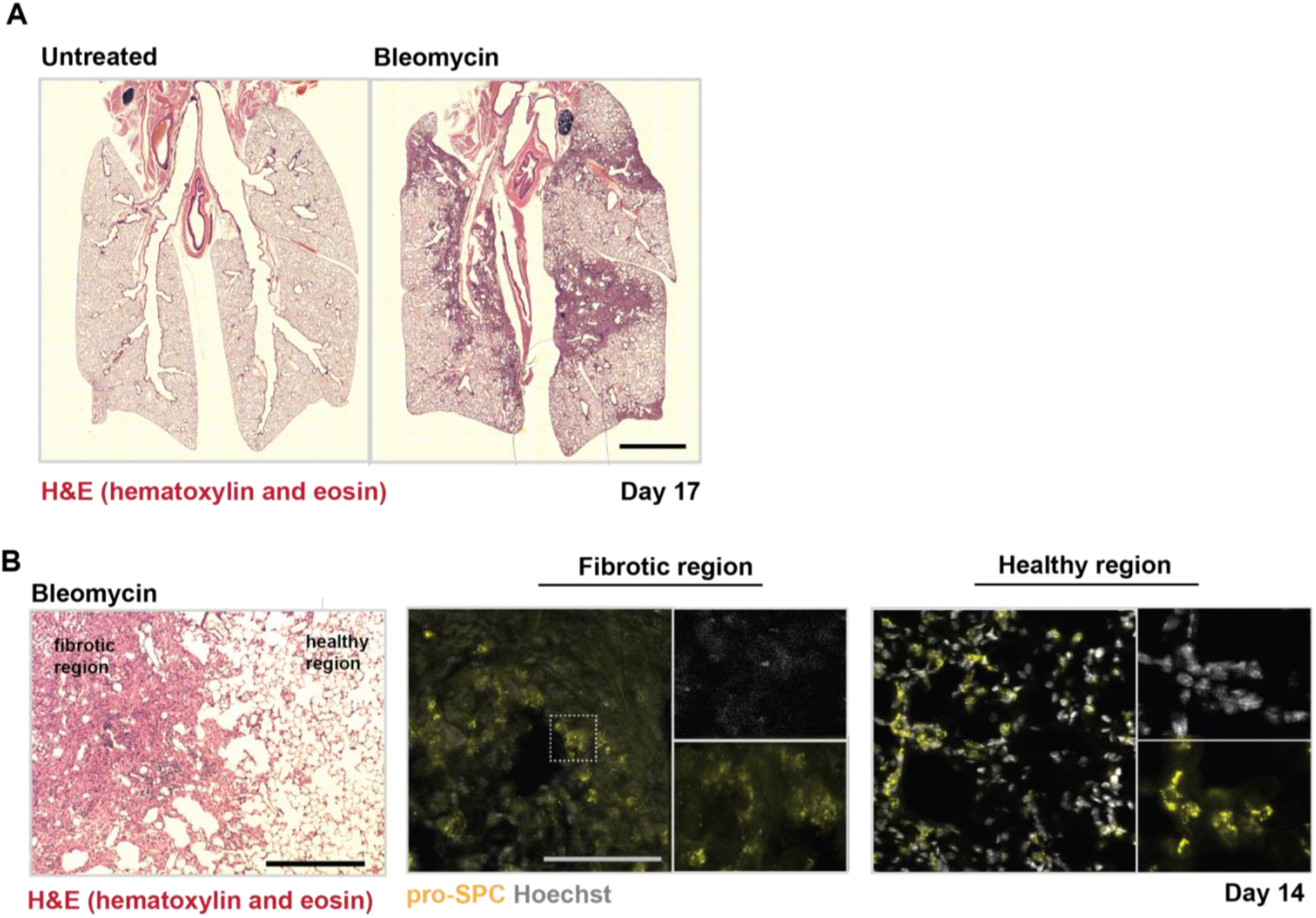
Bleomycin injury of immunocompromised NOD SCID NSG gamma mice. **A** Representative images of hematoxylin and eosin (H&E) staining of fibrotic and healthy lungs of mice 17 days upon bleomycin injury (scale bar 2500 µm). **B** Representative images of fibrotic and healthy regions (H&E, nuclei (grey) and pro-SPC (yellow), scale bars 500 µm (H&E), 100 µm (pro-SPC)).

**Supplementary Figure 11:**
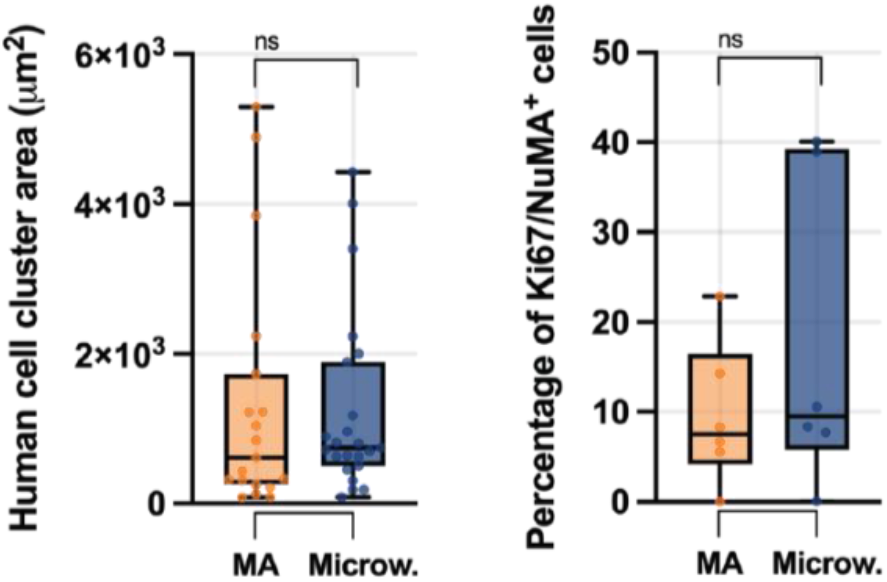
Quantification of transplanted human cell areas. Quantification of the area and percentage of proliferating cells within transplanted human cell area at day 14 (ns = not significantly different by unpaired two-tailed *t*-test, averaged from 3 mice per group, n = 22 lung sections (Matrigel, area), n = 25 lung sections (microwells, area), n = 6 lung sections (Matrigel, microwells).

**Supplementary Figure 12:**
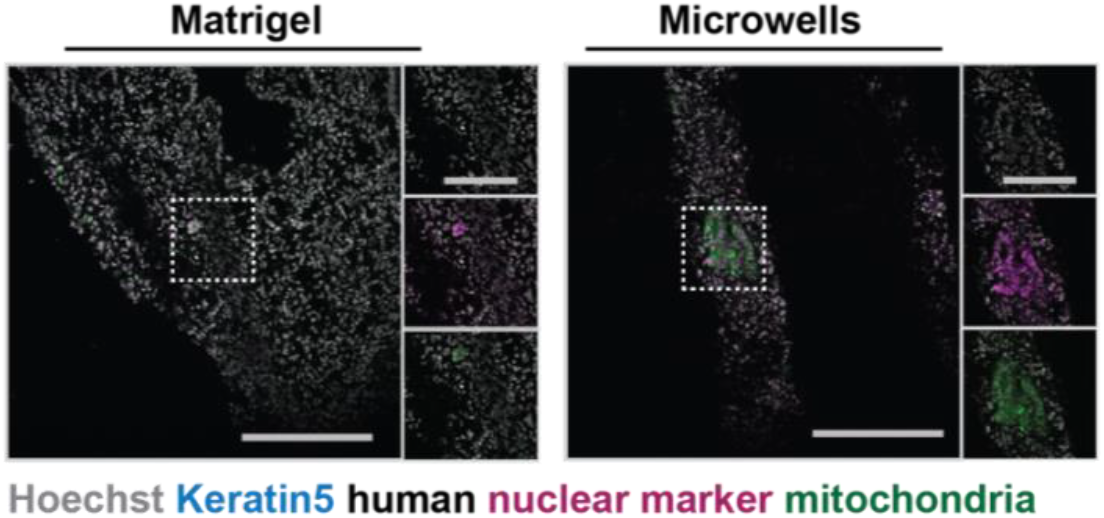
Absence of dysplastic differentiation of human cells upon orthotopic transplantation. Representative images of Keratin 5 expression of transplanted human cells derived from alveolospheres upon culture in Matrigel or atop microwells (nuclei (grey), keratin 5 (cyan), human nuclear marker (magenta), and human mitochondrial marker (green), scale bars 100 µm).

**Supplementary Figure 13:**
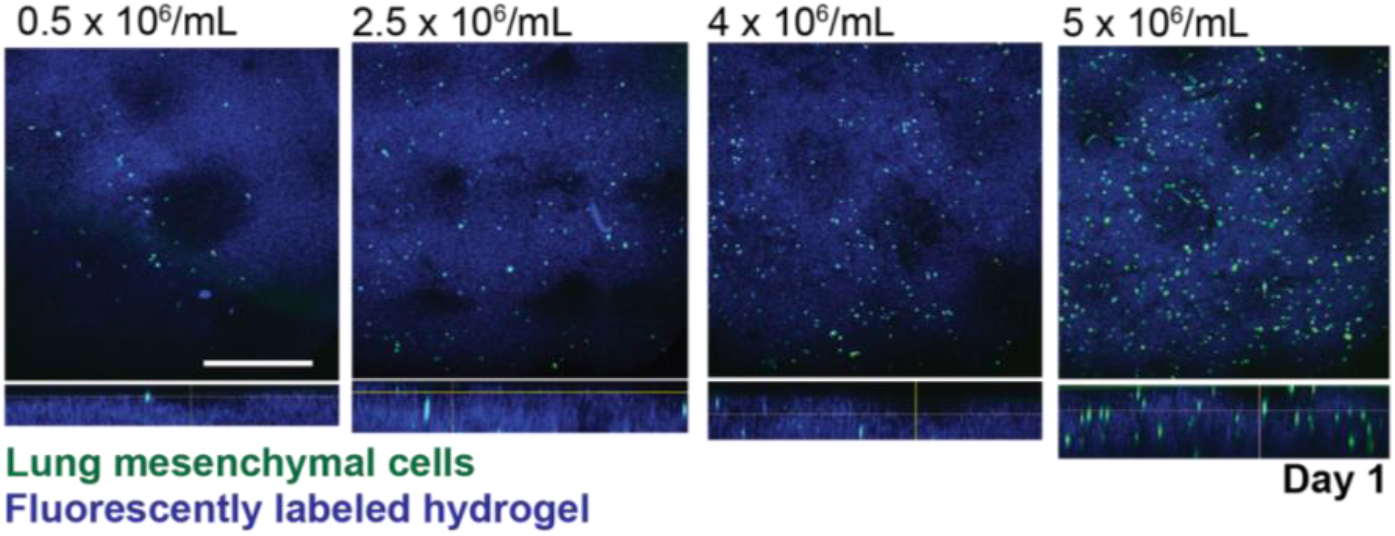
Density of hydrogel embedded lung mesenchymal cells. **A** Representative images of microwell hyaluronic acid hydrogels modified with fluoresceine and various densities of embedded human lung fibroblasts at day 1 of swelling in saline (medium: 500 µm/200 µm (width/depth), cell tracker (green), hydrogel (blue), scale bar 500 µm).

**Supplementary Figure 14:**
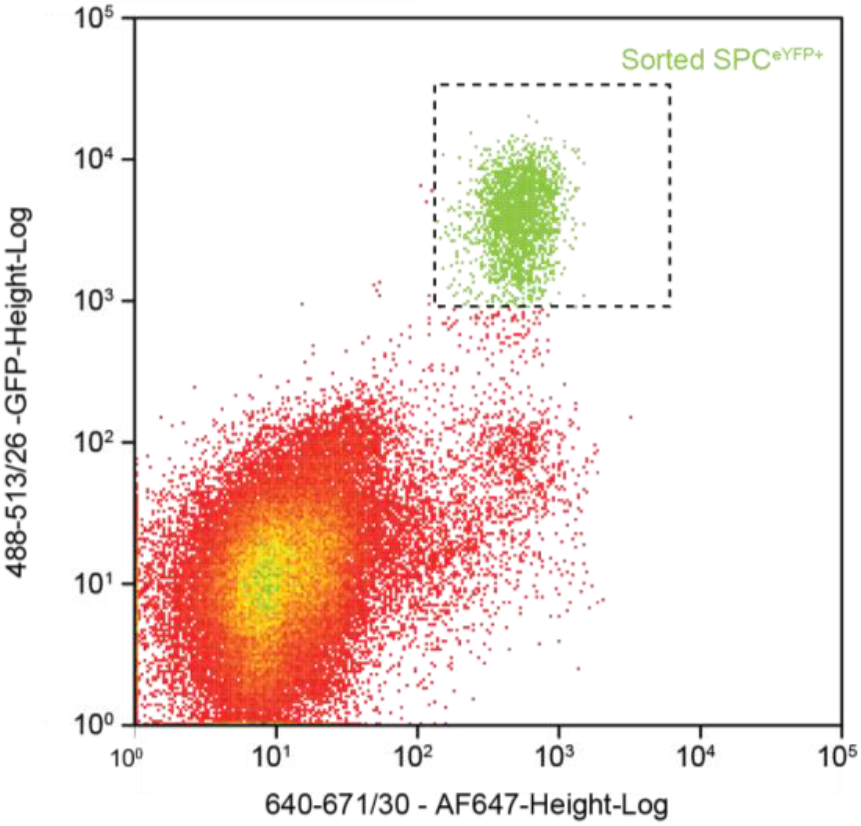
Gating and sorting of primary mouse SFTPC^eYFP+^. Representative flow cytometry plot of cells isolated from SFTPC^CreERT2^; R26R^eYFP^ mice for microwell culture.

**Supplementary Figure 15:**
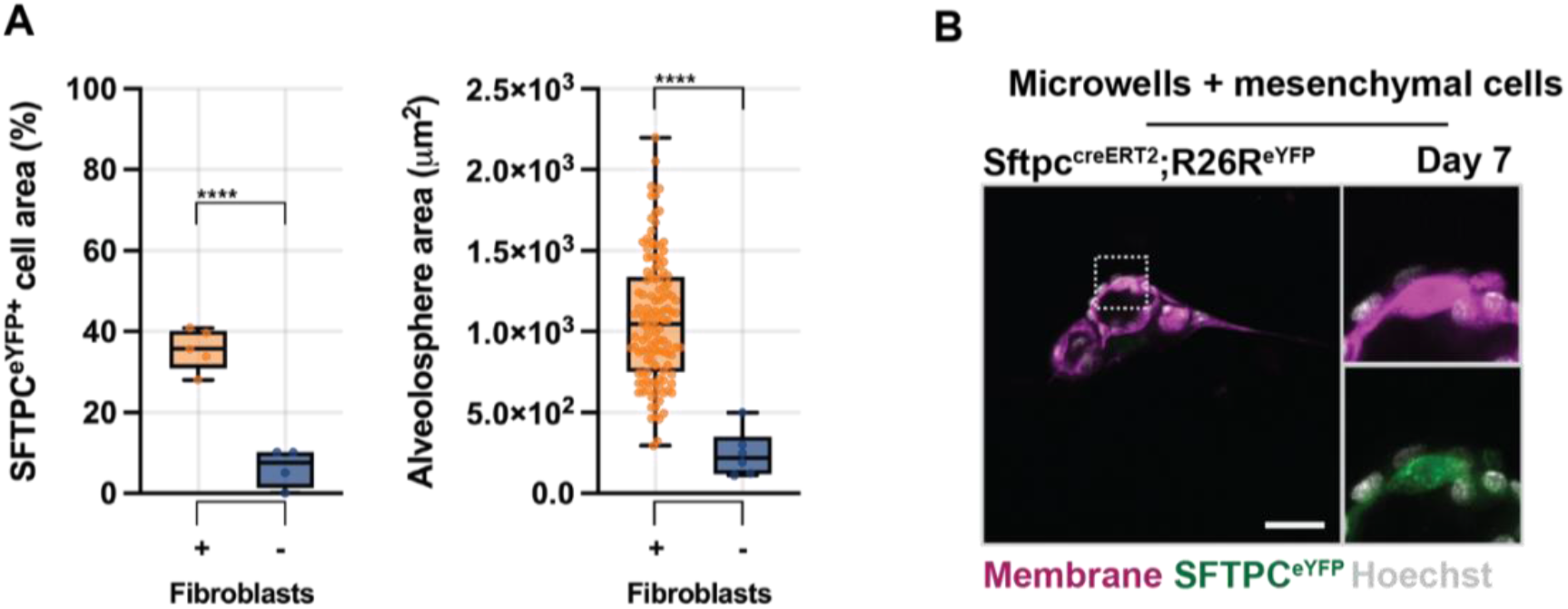
Influence of fibroblast co-culture on SFTPC^eYFP+^ organoid formation within microwell. **A** Quantification of SFTPC^eYFP+^ expression per cell area (cell membrane stain) and area of the projected organoids formed atop of microwells with and without encapsulated fibroblasts at day 7(****p<0.0001 by unpaired two-tailed *t*-test, n = 120 organoids (+) and 10 organoids (-)). **B** Representative image of alveolosphere upon direct co-culture of iAT2 with fibroblasts within microwells at 7 days (mixed in a ratio of 1 to 10, cell mask membrane stain (magenta) nuclei (grey) and SFTPC^eYFP+^ expression (green), scale bar 100 µm).

**Supplementary Figure 16:**
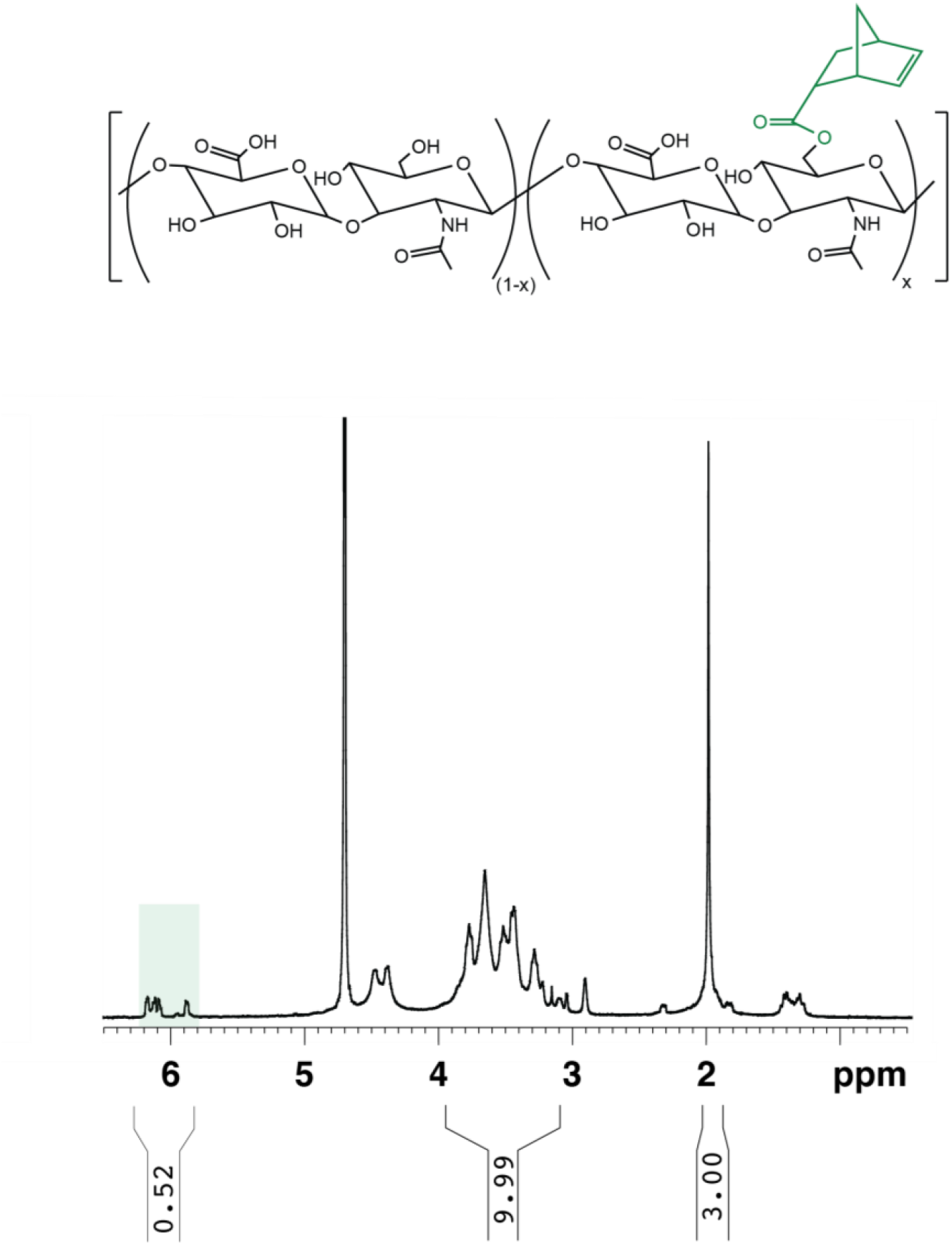
^1^H NMR spectrum of norbornene-functionalized hyaluronic acid (NorHA) in D_2_O. Modification of HA with norbornene (23%) determined by integration of vinyl protons (2H, shaded violet) relative to the sugar ring of HA (10H, shaded grey), analysis was done on each batch used to ensure consistency of modification.

